# Change-point oddball elicits pupil-linked signatures of Bayesian belief updating

**DOI:** 10.64898/2026.09.25.754382

**Authors:** L. Kopel, J.M. Hoomans, E. van Gastel, J.V. Tijhuis, M. van de Graaf, M. Shafik, S. Matthijsse, P.R. Murphy, S. van Gaal, M. Suzuki, J.W. de Gee

## Abstract

The auditory oddball protocol is widely used to study neural responses to unexpected events. In conventional designs, however, stimulus statistics are stationary: deviants do not indicate environmental change and therefore should not drive learning. Here, we introduce “change-point oddball”, a passive, no-report protocol in which the standard-oddball mapping reverses at unpredictable change points, requiring inference over latent environmental states. We measured pupil-linked arousal in three human cohorts. Pupil responses were larger for state-specific oddballs than standards, decayed across newly established standards after state changes, and scaled with model-derived change-point probability. A many-standards control experiment implicated a prediction-based rather than repetition-based mechanism. Finally, within participants, stronger pupil encoding of change-point probability during the passive task was associated with greater change-point-sensitive belief updating in a separate active navigation task. Thus, passive change-point oddball preserves core signatures of adaptive belief updating, enabling cross-species and clinical investigations.

## Introduction

The auditory oddball protocol is widely used in cognitive, systems and clinical neuroscience. In this design, infrequent “oddball” (deviant) stimuli are presented among frequent “standard” stimuli to study how the brain responds to unexpected events [1–4]. A classic physiological correlate of automatic deviance detection is the mismatch negativity, measured with EEG: a negative-going deflection in the difference wave (oddball minus standard) which is maximal at fronto-central scalp sites [1,5–7]. Next to the release of bottom-up neuronal adaptation [8–11], a significant component of the mismatch negativity reflects (model-based) prediction error [6,7,12–14], defined as the difference between predicted and experienced input. This result helped lay the foundation for the influential predictive processing framework [12,15–19] and has led clinical researchers to use the mismatch negativity routinely to characterize abnormalities in information processing across a wide range of neurological and psychiatric conditions [20–22]. More recently, oddball protocols have been paired with invasive recording techniques in monkeys and rodents to pinpoint the precise neural circuits that implement deviance detection [23–28].

A key limitation of the oddball design is that deviants occur as rare events within an otherwise stable environment. Under these conditions, an unexpected stimulus provides little evidence that the generative structure of the environment has changed [29]: the best prediction for the next stimulus remains the frequent standard. This differs from natural environments, in which unexpected events may indicate a change in a latent state and should therefore alter beliefs about future events [29–32]. For example, one or two disappointing meals at your favourite restaurant may signal that the chef has been replaced, prompting an update in expectations about future meals and perhaps leading you to stop going there. The ability to update beliefs appropriately when prediction errors may reflect either transient noise or genuine environmental change is therefore critical for adaptive behaviour. Disruptions in such belief updating have been implicated in psychosis, anxiety, and mood disorders [33–36]. Because a deviant in the classic oddball never signals environmental change, the paradigm cannot tell us how the brain decides whether a prediction error reflects noise or change, which is precisely the computation on which adaptive belief updating depends.

To address this limitation, we developed a modified version of the conventional auditory oddball protocol termed “change-point oddball”. The design comprises two latent states, each specifying reversed stimulus-probability mappings, and probabilistic transitions between states governed by the “hazard rate” [37,38]. Thus, at unpredictable moments, the previous oddball becomes the new standard, and the previous standard becomes the new oddball. An unexpected stimulus with respect to the currently inferred state can therefore provide evidence for a state change, prompting an update in beliefs about the latent state and subsequent stimulus probabilities. Because state changes occur dynamically within a continuous sequence, change-point oddball permits the study of both deviance detection and trial-by-trial belief updating, formalised as Bayesian inference over latent environmental states. Normative Bayesian accounts specify how the impact of prediction errors on learning should depend on their inferred causes, including the probability that the environment has changed [17,19,39–42].

Change-point oddball can be implemented in a passive context, offering several practical advantages over commonly used active reversal-learning or change-point detection tasks [37,43–51]. First, because participants do not need to make overt decisions or reports, the protocol minimizes neural activity related to decision-making, motor preparation and reporting. It is therefore well-suited to study the neural signatures of belief updating in the absence of overt task demands, analogous to no-report protocols in consciousness research [52]. Second, passive presentation makes the protocol particularly suitable for animal models and clinical populations, as it does not require extensive training, task comprehension or an overt response. Finally, change-point oddball can be readily implemented in laboratories that already use conventional oddball. Despite its simplicity, the stimulus sequences generated by change-point oddball allow for defining computational variables (e.g., “change-point probability” derived from Bayesian belief updating models [38,45]) and relating these to stimulus-by-stimulus physiological responses. Thus, change-point oddball can build a direct bridge between the extensive oddball literature and contemporary work on probabilistic inference [37,43–45,50].

Here, we evaluated the utility of change-point oddball by characterizing pupil-linked arousal responses in healthy human participants and linking these to behaviour in an active change-point navigation task. Although most oddball studies have used EEG to measure deviance detection, pupillometry provides an accessible and reliable complementary measure [53–58]. At constant luminance, pupil size fluctuations are associated with activity across multiple neuromodulatory and autonomic systems [59], including, but not limited to, the locus coeruleus–noradrenaline system [60–65], which is thought to encode and broadcast prediction errors [30,66–71]. Pupil responses also track unexpected events and model-derived measures of belief violation in volatile environments [37,43–46,72–81]. We hypothesized that (i) tone-evoked pupil responses are larger for state-specific oddballs vs standards, (ii) following state changes, pupil responses to the newly established standard gradually decrease, (iii) the pupil mismatch response reflects a (model-based) prediction error, (iv) pupil responses linearly track model-derived change-point probability, and (v) individual differences in pupil encoding of change-point probability during passive change-point oddball are associated with the extent to which belief updating during active navigation is modulated by change-point probability.

## Results

### Pupil responses show state-dependent mismatch effects and adjust after latent-state changes

Twenty participants listened passively to sequences of discrete tones while maintaining fixation (**Fig. 1A**). Each sequence consisted of 600 pure tones (stimulus 1, 1 kHz; stimulus 2, 2 kHz) (**Fig. 1B**), generated by one of two latent states that governed stimulus probability: in state A, the probability of stimulus 1 was 0.9 (standard) and the probability of stimulus 2 was 0.1 (oddball); in state B, this mapping was reversed (**Fig. 1C**). After every stimulus, the latent state changed with a hazard rate of 0.05 (**Fig. 1C**), which made state changes unpredictable while maintaining a geometric distribution of state durations (**Fig. S1A**). The resulting probabilistic sequence implemented a volatile auditory environment consisting of 0.5 s tones separated by 1 s inter-stimulus intervals (**Fig. 1D**). For every stimulus we computed computational variables derived from a Bayesian belief updating model [38] as well as stimulus-evoked pupil response amplitudes (**Fig. 1E**; Methods).

**Figure 1.**
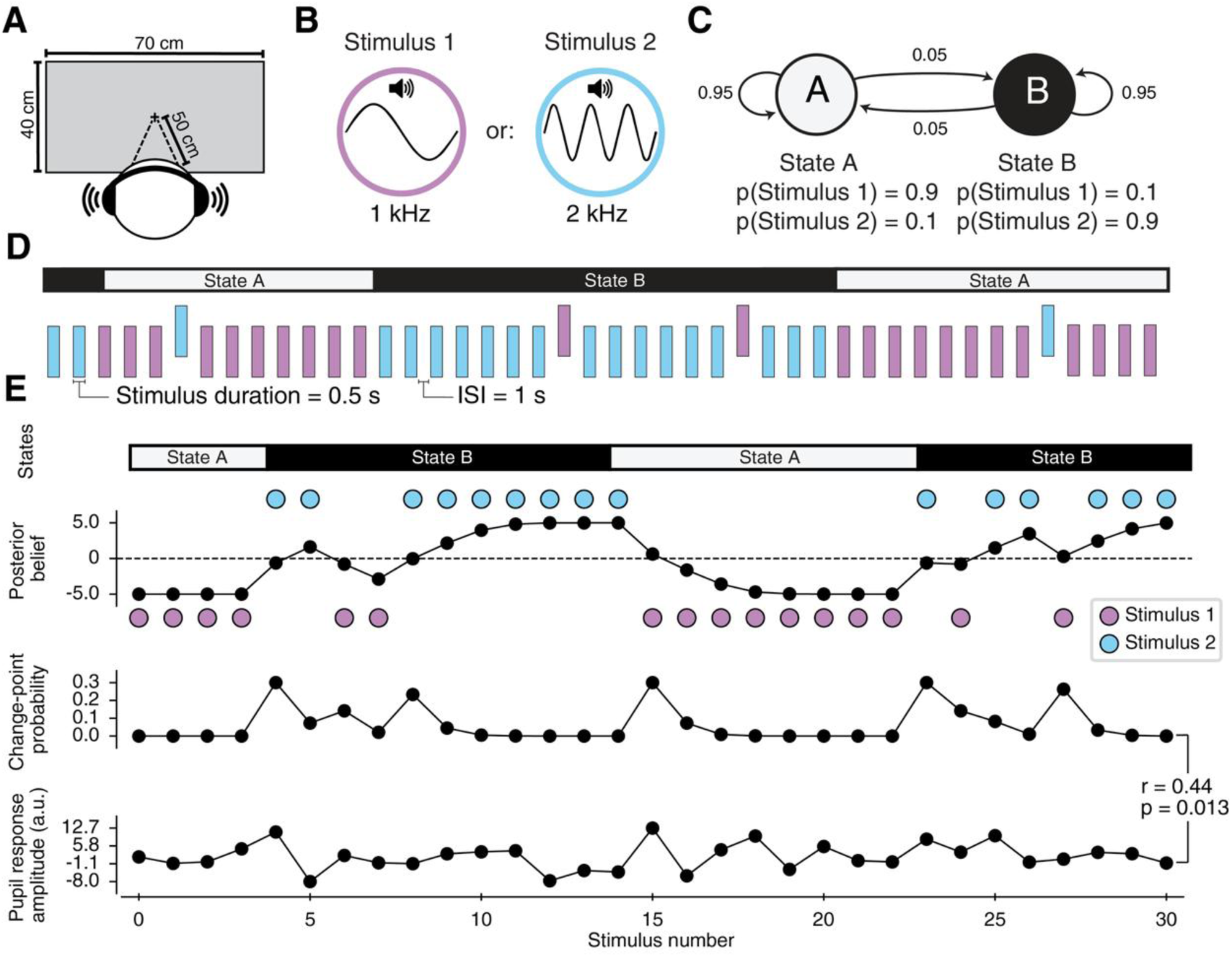
Change-point oddball protocol. **(A)** Experimental setup. **(B)** Auditory stimuli: pure tones of either 1 or 2 kHz. **(C)** Generative process of a sequence. Stimuli are drawn from one of two latent states, which differ in their stimulus-probability mappings: stimuli that are frequent in one state are rare in the other. After each stimulus, the probability of a state change (“hazard rate”) is 0.05. **(D)** Example sequence (segment) of states and stimuli (stimulus duration, 0.5 s; inter-stimulus interval [ISI], 1 s). **(E)** Example sequence (segment) from data set. Posterior belief and change-point probability were derived from a Bayesian belief updating model (Methods; [38]). Pupil response amplitude (Methods) and change-point probability were correlated for this segment. Stats, Pearson’s correlation.

We first tested the hypothesis that stimulus-evoked pupil responses are larger for state-specific oddballs than for the corresponding standards. We therefore predicted an interaction between stimulus identity and latent state. To test this hypothesis, we excluded the first two stimuli following each state change (**Fig. 2A**; 10% of data), when participants’ beliefs about the currently active state were expected to be most uncertain. We first plotted pupil-derivative time courses, indexing the rate of change in pupil size, separately for each stimulus-state combination (**Fig. 2B**). We used the derivative to increase temporal precision [45,82] and specificity for noradrenaline release [65]. Baselined-corrected pupil size time courses are shown in **Fig. S2A**. State-specific oddballs vs standards evoked a larger biphasic derivative response, comprising faster dilation followed by faster constriction (**Fig. 2B**). We next reduced the stimulus-wise pupil-derivative time courses to scalar amplitude measures (Methods) and tested the predicted stimulus-by-state interaction. Specifically, we calculated the difference of differences, [state A stimulus 2 − state A stimulus 1] – [state B stimulus 2 − state B stimulus 1], and tested this quantity against zero (Methods; **Fig. S2C**). As hypothesized, this interaction was significant (**Fig. 2C**), demonstrating that pupil responses to the same physical stimuli depended on the currently active state.

**Figure 2.**
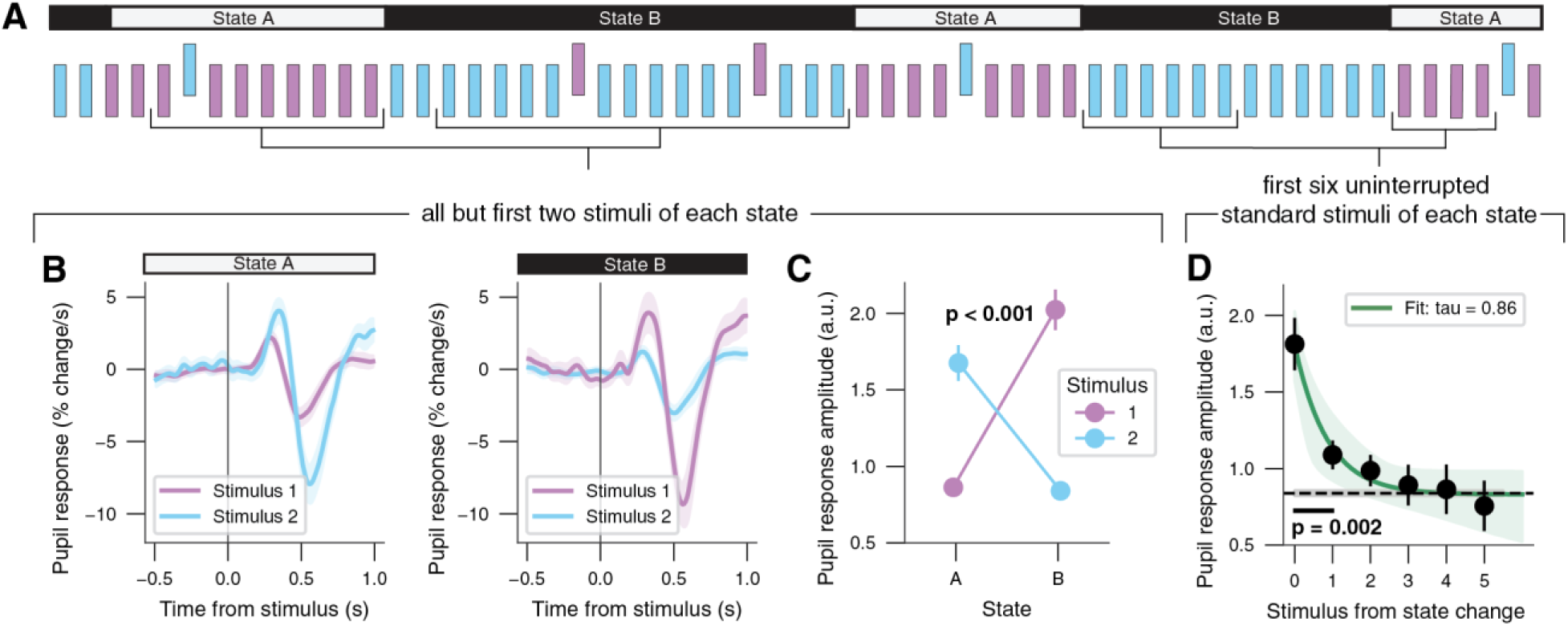
Pupil tracks mismatch responses to state-specific oddballs and adjusts after state changes. **(A)** Example sequence (segment) of states and stimuli. Black brackets indicate stimuli considered for the respective analyses. **(B)** Pupil response time course (first derivative) locked to each stimulus, separately for states and stimuli. Black line, stimulus onset. **(C)** Pupil response amplitude (Methods) separately for states and stimuli. Stats, non-parametric paired permutation test of difference of differences against zero (Methods; see Fig. S2C for permutation null distribution). **(D)** Pupil response amplitude for state-specific standard stimuli locked to state changes (uninterrupted by oddballs; see Fig. S2D for associated pupil response time courses). Green line, exponential decay fit; green shading, 95% bootstrapped confidence interval of that fit; dashed black line, pupil response amplitude to state-specific standards across all stimuli >6 from state change. Stats, cluster-based paired permutation test between data points and dashed line. Panels B-D: shading (excluding green shading in D) or error bars, SEM across participants (N = 20; experiment 1).

This state dependence is consistent with pupil responses adjusting after state changes. Because the probability of an oddball within a state was higher than the probability of a state change (0.1 vs 0.05), the first newly established standard following a state change was initially more likely to be interpreted as an oddball than as evidence for a new state. Accordingly, we next tested the hypothesis that pupil responses to newly established standards gradually decrease following state changes and characterised the timescale of this adjustment. As predicted, the first standard stimulus in a new state evoked a large pupil response, which robustly decreased across subsequent standards in that state (**Fig. 2D**, **Fig. S2C**). The fitted decay time constant was *τ* = 0.86 standards. Thus, after 5*τ* = 5 × 0.86 = 4.3 additional state-specific standards, more than 99% of the initial excess response had decayed. Pupil responses to the first two standards following a state change differed significantly from the average response to stable state-specific standards, defined as the standards occurring at least 6 stimuli after the previous state change (**Fig. 2D**). Thus, pupil-linked arousal declined over the course of repeated (initially surprising) standard stimuli.

Several control analyses speak to the robustness of these results. First, the mismatch response (state-specific oddball vs state-specific standard) was significant when computed separately for each state (**Fig. S2E**). Second, the result does not depend on stimulus selection criteria. In the main analysis, we excluded the first two stimuli after each state change, but the result remains qualitatively the same for different numbers of stimuli from state change as well as for a wide range of previous state durations (**Fig. S2F**). Third, the result does not depend on methodological choices to compute stimulus-wise pupil response scalar amplitudes. In the main analysis, we did so via linear projection [83] of all pupil derivative samples in the range 0-1 s from stimulus onset, including the dilation as well as constriction phase (Methods). The result remains qualitatively the same when isolating the dilation phase (**Fig. S2G**,**H**) or constriction phase (**Fig. S2G**,**I**) with an alternative window-based approach (Methods). Interestingly, the interaction was most pronounced during the constriction phase (**Fig. S2G-I**), which can also be observed at the level of the pupil time courses (**Fig. 2B**). Fourth, we replicated the result in a visual version of change-point oddball (**Fig. S2J**,**K)**, although contrast gratings evoke stimulus-driven pupil constrictions that complicate its combination with pupillometry (**Fig. S2L**).

Finally, we characterized the stability of our effects over shorter and longer time-scales – in other words, how pupil mismatch responses developed over the course of the experiment. Notably, the pupil response interaction was not different between early and late stimuli within a sequence of 600 (**Fig. S2M**; Methods), and, likewise, was not different between early and late sequences (**Fig. S2N;** Methods). The pupil response adjustment after state changes was also not different between early and late stimuli in each sequence (**Fig. S2O**) but was faster in late sequences (**Fig. S2P**). These results indicate that deviance detection and internal belief updating during change-point oddball do not require sustained attention, and that the speed of belief updating may increase after exposure to several sequences.

In sum, pupil responses were larger for state-specific oddballs than for state-specific standards. Following state changes, pupil responses to newly established standards were initially large but declined across several subsequent presentations. These results confirmed our first two hypotheses. This pattern of results may reflect non-mutually exclusive contributions of (i) attenuated responses to repeated standards, including (bottom-up) adaptation or repetition suppression, and (ii) enhanced responses to oddballs that violate (top-down) model-based predictions. In the following analyses, we sought to distinguish between these accounts.

### Pupil mismatch responses are primarily explained by prediction-related rather than repetition-related effects

We next tested the hypothesis that the pupil mismatch response to state-specific oddballs primarily reflects prediction-related rather than repetition-related effects. A new, independent group of fifteen participants completed the change-point oddball protocol and an equiprobable many-standards control experiment [84–86] (Methods). In the control experiment, participants listened to a random sequence comprised of ten distinct tones, each occurring with equal probability (**Fig. 3A**). The 1 and 2 kHz tones used in the change-point oddball protocol were included amongst these ten tones. Because each control tone occurred with probability 0.1, the control experiment matched the probability of state-specific oddballs while avoiding the repeated presentation of a single high-probability standard. Under the conventional logic of the many-standards control, the difference between responses to state-specific standards and control stimuli estimates repetition-related effects, whereas the difference between responses to state-specific oddballs and control stimuli estimates prediction-related effects.

**Figure 3.**
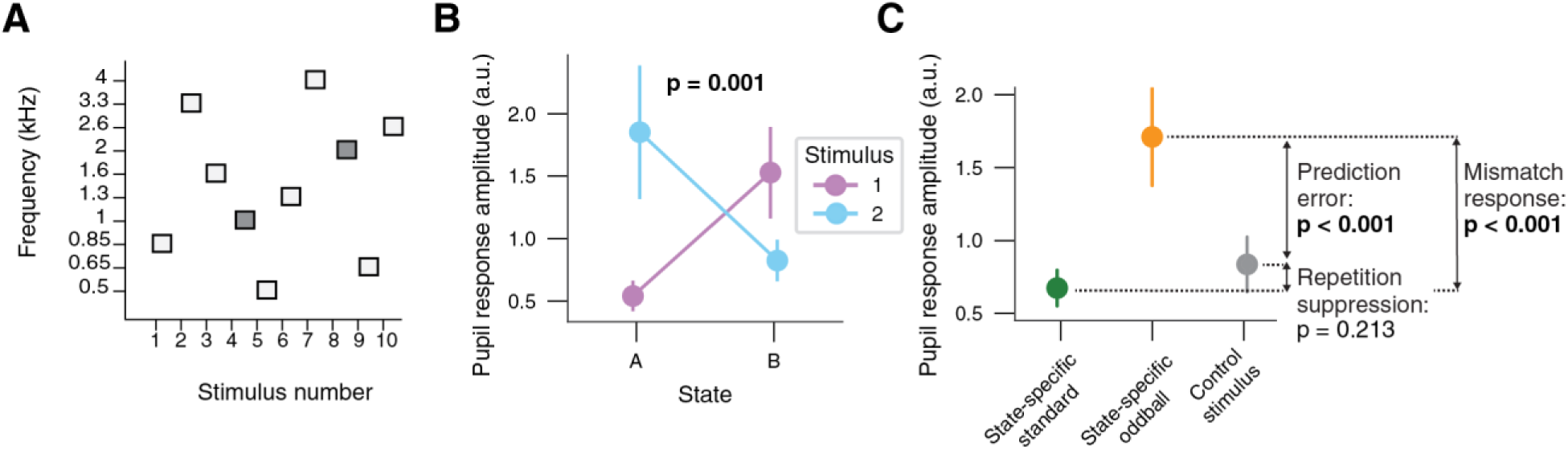
Pupil mismatch response primarily reflects prediction error. **(A)** Example sequence (segment) of many standards control experiment (Methods). Dark grey, frequencies used in change-point odd-ball. **(B)** Pupil response amplitude separately for states and stimuli (for all but first two stimuli of each state). Stats, non-parametric paired permutation test of difference of differences against zero. **(C)** Pupil response amplitude separately for state-specific standards and oddballs in change-point oddball (for all but first two stimuli of each state) and control stimuli (dark grey stimuli in panel A). Stats, non-parametric paired permutation test. See Fig. S3D for pupil response time courses. Panels B, C: error bars, SEM across participants (N = 15; experiment 2).

During change-point oddball, we again observed a significant stimulus-by-state interaction (**Fig. 3B**), replicating the result from the first experiment in an independent sample (**Fig. 2C**). We next recoded the stimulus-by-state combinations as state-specific standards and oddballs (green and orange in **Fig. 3C**) and compared their pupil response amplitudes with responses to the same tones in the equiprobable control context (grey in **Fig. 3C**). Pupil responses to state-specific oddballs were significantly larger than responses to control stimuli (**Fig. 3C**). At the group level, the oddball-control difference accounted for 84% of the total oddball-standard mismatch response. Analyses that separately isolated the dilation and constriction phases yielded qualitatively similar results, although the oddball–control difference was smaller for the dilation phase (**Fig. S3E-I**).

Together, these results suggest that pupil mismatch responses during change-point oddball is driven predominantly by prediction-related effects, with a smaller contribution from repetition-related effects. This finding supports our third hypothesis and is consistent with evidence from conventional oddball protocols [86] and predictive processing accounts of deviance detection [12,15–19].

### Pupil responses track model-derived change-point probability

During active inference tasks, pupil responses track model-derived measures of belief violation [37,43–46,81]. Because passive change-point oddball was designed to probe related computations without requiring an overt response, we tested whether pupil responses also track model-derived change-point probability in this context.

We adopted a widely used Bayesian belief updating model [38,45], which formalises how observers should combine prior beliefs and incoming evidence when latent environmental states can change [87,88]. In this model, the evidence provided by each stimulus is combined with a prior to yield a posterior belief (Fig. 4A). The posterior belief becomes the prior for the next stimulus, after accounting for the overall probability that the latent state changes from one stimulus to the next, called the subjective hazard rate. A subjective hazard rate of 0 yields an identity mapping, corresponding to perfect accumulation: beliefs can become very strong, making the system relatively robust to noise but slow to detect state changes. Increasing the subjective hazard rate (up to 0.5) pulls beliefs towards equal probability of the two states, limiting the accumulation of strong beliefs. This makes the system more sensitive to recent observations but quicker to detect latent state changes.

**Figure 4.**
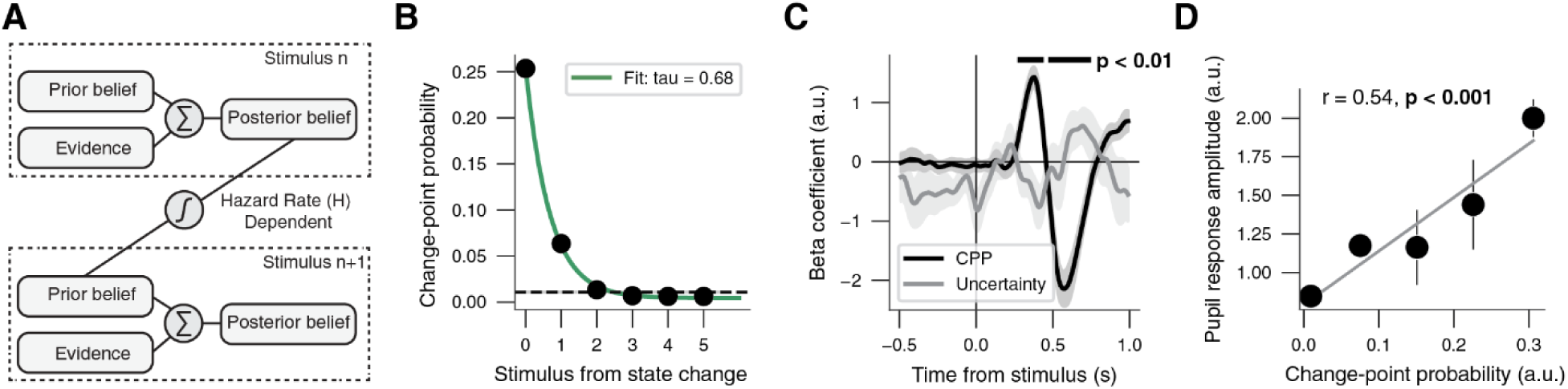
Pupil responses in change-point oddball track change-point probability. **(A)** Schematic illustration of the Bayesian belief updating model (Methods). Evidence provided by each stimulus is combined with a prior belief to yield a posterior belief, which becomes the prior for the next stimulus, after first passing through a non-linear transfer function. **(B)** Change-point probability for state-specific standard stimuli locked to state changes (uninterrupted by oddballs). Green line, exponential decay fit; dashed grey line, change-point probability to state-specific standards across all stimuli >6 from state change. **(C)** Time course of the regression beta coefficients linking pupil response to change-point probability (black) and uncertainty (grey) (Methods). Black bars, timepoints significant change-point probability encoding (cluster-based paired permutation test between beta coefficients and zero vector, first cluster: p = 0.002, second cluster: p = 0.001). **(D)** Correlation between pupil response amplitude and binned change-point probability (see Fig. S4A for histogram and bin edges; Methods). Stats, Pearson’s correlation per participants and non-parametric paired permutation test of r-values against zero vector. Panels C, D: shading or error bars, SEM across participants (N = 20; experiment 1).

The model can be used to compute change-point probability [45], the posterior probability that a change in the latent state has just occurred, given the subjective hazard rate, the evidence carried by a given stimulus and the observer’s belief before encountering that stimulus (Methods). In a change-point oddball sequence (with two distinct discrete stimuli), the change-point probability can be high only when both of the following hold: (i) the observer’s current belief state is strong, and (ii) a new stimulus is inconsistent with this belief.

As predicted, pupil responses during change-point oddball reflected change-point probability. We computed change-point probability for each stimulus in the empirical sequences, using a subjective hazard rate of 0.05 (matching the generative hazard rate; Methods). Strikingly, after state changes, the decay in model-derived change-point probability (Fig. 4B) closely resembled that of pupil response amplitudes (Fig. 2D): the time constant of this decay was *τ* = 0.68, which means that after approximately 5*τ* = 5 × 0.68 = 3.4 additional state-specific standards, more than 99% of the initial excess change-point probability above its asymptote had decayed.

In complementary analyses, we examined the association between pupil response amplitudes and change-point probability across all stimuli, independent of their position relative to state changes. First, we implemented a similar time-wise regression analysis as presented before [45,89]: pupil response predicted by current change-point probability, change-point probability on the preceding stimulus, uncertainty, and baseline pupil size (Methods). We observed robust change-point probability encoding during both the dilation and constriction phases of the stimulus-evoked pupil response (Fig. 4C). Next, using our pupil response scalars, we verified that this effect was indeed linear. To do so, we visualized the distribution of change-point probability (**Fig. S4A**) and binned change-point probability accordingly. Bin-wise pupil response amplitudes increased approximately linearly with bin-wise change-point probability estimates (Fig. 4D). This correlation did not differ significantly between early and late stimuli in the sequence (**Fig. S4B**) or between early and late sequences (**Fig. S4C**).

Finally, we conducted an exploratory analysis to determine which subjective hazard rate produced model dynamics most like the observed post-change pupil response decay. To do so, we computed the change-point probability decay for subjective hazard rates of 0.01, 0.025, 0.05, 0.1, 0.25, 0.5 and 0.75 (**Fig. S4D**). A subjective hazard rate of 0.025 resulted in *τ* = 0.87, which most closely matched the time constant of pupil response amplitude decay after state changes (Fig. 2D). Thus, the temporal dynamics of pupil responses were most consistent with a model that assumed a somewhat more stable environment than the one participants experienced.

In sum, phasic pupil-linked arousal during passive change-point oddball tracked model-derived change-point probability, in line with observations during active inference tasks [45,49]. This confirmed our fourth hypothesis and is consistent with the view that pupil-linked arousal reflects computations involved in Bayesian belief updating.

### Change-point probability encoding in pupil responses during passive change-point oddball is associated with change-point-sensitive belief updating during active navigation

Having established that pupil responses during passive change-point oddball encode model-derived change-point probability, we finally asked whether individual differences in this physiological signature of belief updating were related to an analogous behavioural signature during active inference. To link belief updating across tasks within the same participants, we used change-point probability as a common computational anchor.

A new, independent group of thirty participants completed an audio-visual version of the passive change-point oddball protocol and a virtual-reality change-point navigation task (Methods). During the navigation task, participants were automatically conveyed forward along a virtual track at a constant speed, while controlling their position along the horizontal axis with a joystick. At unpredictable moments, a wall appeared across the full width of the track, containing a gap through which participants had to navigate (Fig. 5A). The horizontal position of this gap was sampled from one of two partially overlapping distributions, corresponding to the latent states (Fig. 5B). Participants moved horizontally to pass through the gap. Successful passage or collision with the wall were accompanied with distinct auditory feedback (Methods). After each wall, the probability of a state change was either 0.08 or 0.12 (Fig. 5C). This hazard rate alternated across blocks (Fig. 5D). Because we found no evidence that behaviour differed between hazard rate conditions (**Fig. S5A,B**), the primary analyses were collapsed across conditions. We therefore used the average generative hazard rate of 0.1 to compute model-derived variables; we additionally report analyses using condition-specific hazard rates (**Fig. S5O-R**).

**Figure. 5.**
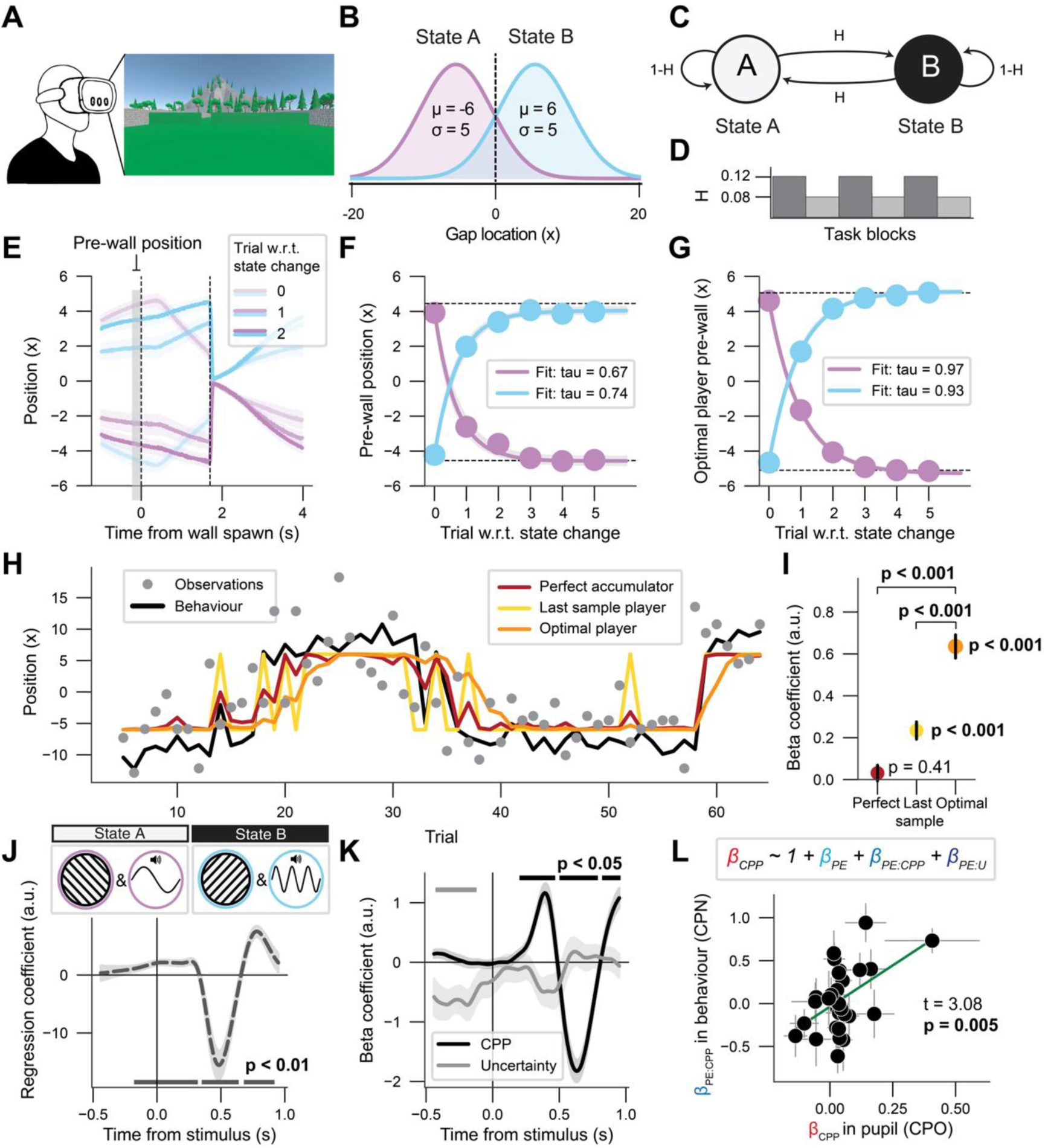
Change-point probability encoding in pupil responses during passive oddball is associated with change-point-sensitive belief updating during active navigation. **(A)** Experimental set-up of the change-point navigation task in virtual reality. **(B)** Two partially overlapping distributions of gap locations implemented the latent states. **(C)** Generative process of a sequence. H, hazard rate. **(D)** Block-wise hazard rate was 0.08 or 0.12. **(E)** Horizontal position aligned to state changes, separately for states and the first three trials with respect to state changes and collapsed across hazard rates. Grey rectangle, time window to calculate pre-wall position. **(F)** Pre-wall position locked to state changes, separately for states and collapsed across hazard rates. Pink and blue lines, exponential fits; light grey shading, 95% bootstrapped confidence interval of fits; dashed black lines, pre-wall position across all stimuli >5 from state change. **(G)** As F, but for optimal Bayesian player (Methods). **(H)** Example sequence (segment) from data set. Grey dots and black line indicate observations (gap locations) and pre-wall position behaviour of example participant. Coloured lines represent pre-wall positions of three synthetic players (Methods). **(I)** Group-level multiple linear regression coefficients from modelling empirical pre-wall position as function of positions of the different synthetic players (Methods). Stats, non-parametric paired permutation test against zero. **(J)** Top: Audio-visual change-point oddball protocol: 1 and 2 kHz tones paired with -45° and 45° oriented Gabor patches, respectively. Bottom: intercept time course of the time-wise regression of pupil derivative on computational variables (in panel K; Methods). Bars, significant timepoints (cluster-based paired permutation test between intercept and zero; all clusters, p < 0.01). **(K)** Time course change-point probability encoding (black) and uncertainty encoding (grey; Methods). Black bars, significant timepoints (cluster-based paired permutation test between beta coefficients and zero; p = 0.001, p = 0.034 and p = 0.001 for the three clusters, respectively). Grey bar, as black bar (p = 0.017). **(L)** Change-point-sensitive belief updating in change-point navigation task (βPE:CPP) plotted against change-point probability encoding in pupil during passive change-point oddball (βCPP). Every data point is a participant. Stats, across-subjects multiple linear regression (Methods); error bars, bootstrapped 95% confidence interval (10K bootstraps; Methods). Panels E, I, J, K: error bars, shading, SEM across participants. All panels: N = 30 (experiment 3).

Crucially, the task timing encouraged participants to position themselves before each wall appeared (“wall spawn”). After each wall, participants were automatically returned to the centre of the track, such that pre-wall position reflected their prior belief about the location of the upcoming gap rather than their position on the preceding trial. Participants adjusted their pre-wall positions according to the currently active latent state and updated these positions following state changes (Fig. 5E**,F**). We quantified pre-wall position as the average horizontal position from -0.2 to 0 s before wall onset. Following state changes, pre-wall position gradually adjusted towards the location appropriate for the new state (Fig. 5F). The fitted time constants were τ = 0.67 trials for transitions from state A to B, and τ = 0.74 trials for transitions from state B to A. Thus, after approximately 5τ, or 3-4 additional trials, more than 99% of the initial adjustment in pre-wall position had occurred.

We next asked which belief-updating strategy best captured participants’ pre-wall positions. We compared their behaviour with that of three synthetic players (Methods): (i) an optimal player, which integrates gap positions according to a normative Bayesian change-point model with a subjective hazard rate matching the generative task statistics; (ii) a last-sample player, which positions itself according to the location of the gap on the preceding trial only; and (iii) a perfect accumulator, which integrates all previous gap positions without allowing for state changes, corresponding to a subjective hazard rate of 0. At the group level, participants’ behaviour was best captured by the optimal player (Fig. 5G**–I**). First, the optimal player time course of adjustment in pre-wall position following state changes (Fig. 5G) most closely resembled the empirical time course (Fig. 5F); updating in the last sample player and perfect accumulator was substantially faster and slower, respectively (**Fig. S5C,D**). Second, the mean absolute error between empirical and simulated pre-wall positions was smallest for the optimal player for 19 out of thirty participants (Fig. 5H**, Fig. S5E**). Third, across participants, empirical pre-wall positions were more strongly predicted by those generated by the optimal player than by positions generated by either alternative player (Fig. 5I). Together, these results indicate that participants’ behaviour was best described by a normative belief-updating strategy that accounts for the possibility of latent-state changes.

Having established this group-level correspondence with normative belief updating, we next quantified individual differences in change-point-sensitive updating. We defined the belief update on trial *t* as the change in pre-wall position from trial *t* to *t+1* and modelled this quantity using a subject-wise linear regression [50,90] as function of (i) the prediction error (β_PE_), (ii) the interaction between prediction error and change-point probability (β_PE*CPP_), and (iii) the interaction between prediction error and uncertainty (β_PE*U_; Methods). The prediction error was defined as the trial-wise difference between pre-wall position and gap location (Methods). The prediction-error-by-change-point-probability interaction quantified the extent to which each participant increased belief updating when a prediction error was more likely to signal a state change. Most participants showed a significant modulation of prediction-error-based updating by change-point probability, although there were substantial individual differences (**Fig. S5F**). These participant-wise behavioural signatures of adaptive belief updating showed high test-retest reliability (r = 0.89; **Fig. S5G**,**H**), allowing us to test its association with pupil responses during passive change-point oddball.

Before the navigation task, the same group of thirty participants was naively exposed to an audio-visual version of the passive change-point oddball protocol (Fig. 5J). We replicated the pupillary stimulus-by-state interaction (**Fig. S5I**), although it was likely attenuated by contrast-driven pupil constriction (**Fig. S5J**). The same time-wise regression as before (Fig. 4C) captured this constriction component and revealed robust change-point probability encoding (Fig. 5J,**K**). This participant-wise physiological signature of adaptive belief updating showed high test-retest reliability (r = 0.91; **Fig. S5K**,**L**). In a multiple linear across-subjects regression model, we used the individual regression coefficients β_PE_, β_PE*CPP_, β_PE*U_ from the change-point navigation task (described above) to predict change-point probability encoding in the pupil during passive change-point oddball. As predicted, the strength of change-point probability encoding in the pupil was predicted by strength of change-point probability modulation of belief updates in the change-point navigation task (β_PE*CPP_) (Fig. 5L), but not by strength of uncertainty modulation (β_PE*U_) or prediction error on belief updates alone (β_PE_) (**Fig. S5M**,**N**). Thus, people with higher change-point probability encoding in the pupil during passive change-point oddball updated their beliefs more after likely change points in an active change-point navigation task.

These results are robust to the specifics of the analysis. We repeated the regression with the pupillary stimulus-by-state interaction (**Fig. S5I**) as dependent variable. Again, strength of stimulus-by-state interaction in the pupil during change-point oddball was predicted by strength of change-point probability modulation of belief updates in the change-point navigation task (β_PE*CPP_: t = 3.08, p = 0.005). Repeating both analyses after removing one potential outlier yielded qualitatively similar results: t = 2.06, p = 0.050 and β_PE*CPP_: t = 2.39, p = 0.025, for change-point probability encoding and stimulus-by-state interaction as dependent variable, respectively.

In sum, individual differences in change-point probability encoding in pupil responses during passive change-point oddball were associated with the extent to which participants adjusted their beliefs following prediction errors that likely signalled latent-state changes during active navigation. These findings raise the possibility that partially shared mechanisms support change-point-probability encoding across passive and active contexts and support the idea that passive change-point oddball can be used to study belief updating.

## Discussion

Here we introduce change-point oddball, a passive auditory protocol in which the roles of standard and oddball stimuli reverse at unpredictable moments, such that the state of the world is latent and must be continuously inferred. This provides the opportunity to study not only deviance detection, as in conventional oddball, but also (Bayesian) belief updating after state changes. Consequently, change-point oddball is suited to efficiently study the full cycle of prediction error signalling and belief updating. In our study, human participants were passively exposed to auditory change-point oddball while we simultaneously measured their pupil size at constant luminance. First, we observed multiple pupil-based signatures of belief updating: (i) larger tone-evoked pupil responses for state-specific oddballs vs standards, which primarily reflected (top-down) prediction error rather than the release of (bottom-up) repetition suppression, (ii) exponentially decaying pupil responses to state-specific standards after state changes, and (iii) a linear relationship between pupil responses and change-point probability, derived from a Bayesian belief updating model. Second, we showed that pupil-based signatures of belief updating measured in passive change-point oddball are associated with behavioural signatures of normative belief updating in an active inference task. Thus, phasic pupil-linked arousal may support internal belief updating after unexpected outcomes.

Passive change-point oddball bears a close conceptual relationship to the statistical learning literature but offers several advantages. In one series of experiments, participants heard brief sequences of rapid tone-pips (clip duration, 5–7 s, tone-pip rate, 20 Hz) forming a random or regular pattern that contained a single state change [73,75,79]. Participants were instructed to attend to and report short silent gaps that were uncorrelated to state changes. Regularity violations (regular to random, regular to different regular) but not regularity emergence (random to regular) evoked pupil-linked arousal (unless the regular pattern was simple and made up of only one or two tone-pips). The state changes embedded in change-point oddball constitute precisely this type of abrupt regularity violation, as the previously reliable standard-deviant mapping is suddenly invalidated. In another study, participants passively listened to sequences of tones that were presented at a rate of 4 Hz, with a certain probability to sample frequencies from one of two Gaussian distributions [74]. Here too, the generative process could undergo one state change during a sound clip of 15 s. Pupil dilation scaled with model-derived surprise and precision. Change-point oddball offers a number of advantages compared to both designs discussed here: (i) the 1 s inter-stimulus interval is sufficient to capture individual tone-evoked pupil responses without overlap, eliminating the need for convolutional approaches based on a time-invariant linear model [83,91,92] that relies on assumptions that may not hold in practice [93,94], (ii) related, the ability to characterize the timescale of belief updating (Fig. 2D), (iii) state changes are completely unpredictable instead of always occurring between 2.5 s and 3.5 s [73,75,79], or between 3.75 s and 6.25 s [74], from sound clip start. Furthermore, compared to the rapid tone-pip protocol [73,75,79], change-point oddball allows for the use of Bayesian belief updating models to compute stimulus-wise computational variables (every stimulus is associated with a log-likelihood ratio of being generated by latent state A vs B; see also next paragraph). One downside of the current implementation of change-point oddball is that it generates fewer state changes per unit time (2 per minute vs 3.75 per minute [73,75,79] or 2.9 per minute [74]) and, therefore, longer experiments may be required to reliably characterize the neural circuits that implement belief updating. Future work should try increasing the hazard rate (e.g., to 0.1, which would approximately double the number of state changes per time interval).

Passive change-point oddball also has a close conceptual link to the (Bayesian) belief-updating literature [37,43–45,49,95], while again offering several advantages. In predictive inference tasks, participants integrate sensory information to infer latent states and, on every trial or intermittently, predict the next sample or report on the current state of the environment. Within this framework, evoked pupil responses are typically associated with violations of learned, top-down expectations [37,43–46,49]. Our findings align with this view: using a many-standards control experiment, we showed that pupil mismatch responses during change-point oddball primarily reflect top-down prediction error, and we further demonstrated that pupil responses scale linearly with change-point probability. In addition, we showed that the strength of change-point probability encoding in passive pupil responses is correlated with increased updating after likely change-points in an active inference task. Compared with active inference tasks, passive change-point oddball offers several benefits. First, it does not require behavioural responses on each trial or intermittently, avoiding potential disruptions of the ongoing inference process. Second, neural activity related to decision-making and reporting is reduced, making the protocol well suited to study belief updating in the absence of overt decisions or reports, analogous to no-report protocols in consciousness research [52]. Third, it is particularly suitable for animal models and clinical populations, as it eliminates the need for extensive training and sustained conscious attention. A limitation of change-point oddball is that the protocol uses only two discrete stimuli, rather than sampling from distributions that instantiate the latent states. Consequently, the absolute log likelihood of each stimulus is fixed, which limits the complexity of evidence integration, and modelling thereof, our protocol can support. Future work should address this limitation by implementing a continuous version based on stimulus distributions. Having said that, despite our simplification, we did identify pupillometric signatures of belief updating in a passive setting that were associated with active belief updating behaviour. Thus, passive change-point oddball can be seen as the most basic version of a (perceptual) inference protocol.

Change-point oddball fundamentally differs from flip-flop and local-global designs. In the flip-flop design, the roles of standard and deviant are reversed between blocks, effectively controlling for stimulus-specific effects on mismatch responses [96]. The local global protocol introduces a hierarchical structure by dissociating short-timescale (local) from long-timescale (global) prediction errors, enabling the separation of automatic sensory mismatch from higher-order conscious processing [97]. Critically, in both designs, the statistical structure is stationary within blocks, meaning that participants do not need to continuously update their belief based on the unfolding stimulus sequence to (implicitly) infer latent states. Thus, change-point oddball allows for probing continuous belief updating while flip-flop and local-global designs do not. Change-point oddball is more closely related to the roving oddball design, in which a stimulus is repeated a variable number of times before a different stimulus starts a new train, so that each deviant becomes the next standard [98]. In the roving design, however, every deviant signals a change in the standard. Deviance and change are thus confounded, and observers never need to infer whether a deviant reflects noise or a new state. Computational single-trial analyses of roving and volatile oddball sequences have shown that mismatch responses track model-derived prediction errors and their precision [98–100] dependent on volatility [101–103] . Change-point oddball complements these designs by making the relationship between deviance and change probabilistic: a single state-specific oddball most likely reflects noise, whereas a run of them signals a likely state change. This allows deviance detection and inference about latent-state changes to be studied within one continuous sequence.

Phasic pupil-linked arousal was heighted after unexpected events including state changes, but how does this support internal belief updates? Influential theoretical work assigns a central role to the noradrenergic locus coeruleus (LC): phasic noradrenaline may act as an interrupt signal triggered by unexpected events, resetting ongoing computations and driving updates of the belief state [66,67,69–71,104]. Consistent with this, noradrenaline has been shown to shape learning rates in volatile environments [70,81,105]. One scenario consistent with our data is that phasic noradrenaline release changes the relative strength of information flow between cortical processing stages, suppressing top-down relative to bottom-up signals [19]. According to the influential predictive coding framework, top-down prediction from higher areas in the cortical hierarchy are compared to bottom-up sensory signals from lower cortical areas [17–19,39–42,106]. When these predictions are not met, a prediction error is evoked, which travels up the hierarchy and ultimately acts as a teaching signal in order to update the belief state and refine future prediction accordingly [32,106]. In the perceptual inference process studied here, early sensory cortices provide bottom-up likelihood signals, whereas top-down signals encode prior beliefs [19,107]. Phasic arousal could therefore reduce the weight of the prior relative to the sensory likelihood. An alternative hypothesis follows more recent interpretations of the predictive coding framework, which favours the flexible imprinting of predictions in lower areas by higher areas through the means of oscillations rather than strict prediction error cascades [108]. In this scenario, noradrenaline release supports the preparation of lower areas by shifting gain according to top-down signals. Future work should characterize whether noradrenaline supports the updating of beliefs by shifting the information flow in favour of either (i) bottom-up sensory signals, (ii) top-down model-based instructions or (iii) both [44].

Individual differences in pupil encoding of change-point probability during passive change-point oddball predicted how strongly the same participants increased belief updating after likely change points during active change-point navigation. This is remarkable, because the two tasks share little beyond their computational structure: they differ in sensory format, timescale, hazard rate and the presence of actions and outcome feedback, and they were administered in separate sessions. Moreover, the association was specific to the change-point-dependent modulation of learning (β_PE*CPP_) and was absent for the overall influence of prediction errors (β_PE_) and its modulation by uncertainty (β_PE*U_), which argues against general differences in engagement or pupil responsiveness. We propose that this shared component reflects individual differences in the gain of the phasic locus coeruleus–noradrenaline (LC–NA) response to evidence for a change point. Ithas been proposed that phasic LC activity evoked by unexpected events increases the influence of new observations on beliefs [66,67,69–71]. LC neurons broadcast prediction errors across the cortex to promote plasticity [109], and pharmacological manipulations of noradrenergic transmission alter belief updating and learning in changing environments [81,110,111]. An individual whose LC responds more strongly to change-point evidence would therefore show both larger change-point-related pupil responses during passive listening and stronger change-point-dependent updating whenever beliefs guide behaviour. The individual differences need not originate in the LC itself: the LC receives input from prefrontal cortex [60,112–114], and cortical circuits implicated in computing change-point probability [45,50] may set the strength of its response. In either case, LC–NA would provide a common, domain-general route through which evidence for a change point is converted into learning, although pupil size also reflects other neuromodulatory systems (see below). This hypothesis yields testable predictions. Pharmacological manipulation of noradrenergic activity in humans, or optogenetic manipulation of the LC in animals exposed to change-point oddball, should alter both pupil encoding of change-point probability and subsequent updating. In addition, trial-by-trial change-point-evoked pupil responses during change-point oddball may be a sensitive biomarker for LC-NA structural and functional integrity, for example in age-related cognitive decline or schizophrenia [89,115]. Given the modest sample and significance, and fixed session order (the passive session always came first), the association should also be replicated in larger samples.

Using pupillometry to track deviance detection and belief updating in change-point oddball comes with both advantages and limitations. Compared to EEG, pupillometry is inexpensive, mobile, and non-invasive (does not require skin contact / gel), making it particularly suitable for clinical applications [89,116,117]. Pupillometry is also integrated in most head-fixed monkey and mouse setups [63,118], which makes it an attractive measure for cross-species research [82,119]. Pupil size is however a relatively unspecific measure [59]: at constant luminance, fluctuations reflect activity in the cortical salience network [61] and several subcortical neuromodulatory structures [63], such as the noradrenergic locus coeruleus [61,62,65], the cholinergic basal forebrain [61,65,120], the serotonergic dorsal raphe [64,121] and likely others [61,121,122]. Future studies should therefore isolate the precise neuroanatomical and neurochemical sources of the effects reported here. Pupil size is also strongly driven by low-level visual features such as luminance and contrast [123,124]. Notably, despite the robust contrast-related constrictions (**Fig. S2J**), we still observed mismatch responses and change-point encoding (**Fig. S2K**, Fig. 5J). Nevertheless, recent work indicates that arousal-related pupil responses interact with luminance in a nonlinear, multiplicative manner [93,94]. Future studies aiming to implement visual change-point oddball should therefore use stimuli designed to minimize the pupil light reflex (for example, low-contrast stimuli that primarily vary in spatial features) or complement pupillometry with electrophysiological measures such as EEG.

Taken together, change-point oddball is a powerful and scalable tool for studying prediction-error-driven belief updating across species, recording methods and clinical groups. We advocate for using change-point oddball in study contexts ranging from animal models to clinical populations, to investigate underlying mechanisms and characterize aberrant belief updating.

## Methods

### Participants

Twenty participants (11 females; 18-23 y) participated in experiment 1. Fifteen participants (9 females; 20-25 y) participated in experiment 2. Thirty-one participants (18 females; 18-24 y) participated in experiment 3. All participants were healthy and had normal or corrected-to-normal hearing and vision. Participants gave written informed consent and were remunerated by the hour or received credit points. Experiment 1 was approved by the ethics committee of the Department of Psychology at the University of Amsterdam (FMG-7300), experiments 2 and 3 by the ethical committee for the natural sciences at the University of Amsterdam (FNWI-282 and FNWI-518, respectively). In experiment 3, one participant was excluded from the analyses, because more than 50% of trials had to be excluded due to blinks during passive change-point oddball.

### Procedure

During all change-point oddball sessions, participants were asked to sit in a dimly lit room in front of the screen at 50 cm distance resting their head on a chin rest. Participants were instructed to passively perceive the stimuli and focus their gaze on a black fixation mark placed in the middle of grey background (Fig. 1A), while we recorded pupil size at constant luminance.

#### Experiment 1 (change-point oddball)

In two experimental sessions of two hours each, participants passively perceived six sequences of 600 auditory stimuli, and six sequences of 600 visual stimuli, yielding a total of 3600 auditory stimuli and 3600 visual stimuli. The auditory and visual sequences were presented in alternating order, and which perceptual domain came first was counterbalanced across participants. Each sequence followed a passive oddball design, with stimulus 1 (the “standard”) appearing 90% of the time and stimulus 2 (the “oddball”) appearing 10% of the time (Fig. 1B,**C**). Critically, we additionally introduced (i) two states, defined by their reverse mapping between the two stimuli and their probabilities, and (ii) a 0.05 probability (“hazard rate”) of a state change after every stimulus presentation (Fig. 1C,**D**). This ensured that both stimuli appeared equally often within a sequence, and that oddball status was dependent on the local state (Fig. 1C,**D**). In all cases, the stimulus duration was 0.5 s (including fade in and fade out) and the inter-stimulus-interval was 1 s. In the auditory sequences, stimuli were pure tones (1 kHz and 2 kHz) (Fig. 1B). When functioning as a state-specific oddball, the 1 kHz tone elicited a slightly larger pupil response than the 2 kHz tone (**Fig. S2B**). In the visual sequences, stimuli were full-contrast Gabor patches (radius, 20 degrees visual angle, spatial frequency, 0.8 cycles per degree of visual angle, rotation, -45 or 45 degrees). The 1 kHz tones were presented at 63 dB and the 2 kHz tones at 66 dB (to adjusted for the equal-loudness contour) via an IMG Stageline MD-5000DR over-ear headphone. Visual stimuli were displayed on a 27’’ LG UltraGear monitor operating at 100 Hz.

#### Experiment 2 (many standards control)

In one experimental session of two hours, each participant passively perceived six sequences of 600 auditory stimuli, yielding a total of 3600 auditory stimuli. Three sequences were the same as described under Experiment 1 (1800 change-point oddball stimuli). In the three remaining sequences, 10 different tones (0.5, 0.65, 0.85, 1, 1.3, 1.6, 2, 2.6, 3 and 4 kHz) occurred equally often and were played in random order (1800 “many standards” control stimuli) (Fig. 3A). Again, tone duration was 0.5 s and ISI 1 s. The 1 kHz tones were presented at 63 dB and the 2 kHz tones at 66 dB (to adjusted for the equal-loudness contour) via Logitech Z150 speakers. Visual stimuli (grey background and fixation dot) were displayed on a 32’’ ASUS ROG monitor operating at 100 Hz.

#### Experiment 3, session 1 (audio-visual change-point oddball)

In one experimental session of two hours, each participant passively perceived five sequences of 600 audio-visual stimuli, yielding a total of 3000 audio-visual stimuli. Stimuli were the same as in experiment 1 but combined across modalities with the goal of maximising salience (1 kHz pure tone and -45 degrees Gabor patch or 2 kHz pure tone and 45 degrees Gabor patch; oddball status was the same in both modalities). Stimuli were presented with the same hardware as in experiment 2.

#### Experiment 3, session 2 (change-point navigation)

In one experimental session of one hour and thirty minutes, each participant completed six blocks of 12 minutes (133– 147 trials) of a Virtual Reality (VR) experiment. The task was presented using a Meta Quest 3 headset and implemented in Unity (version 6.2). The VR environment appeared as a green track with stone walls on either side, extending 20 x-units to either side of the middle of the track. Participants moved forward along the track with a constant speed of 4.86 x-units per second and controlled lateral movement via a joystick. At unpredictable moments, a green wall appeared across the full width of the track, containing a gap through which participants had to navigate. The x-position of the gap was sampled from one of two partially overlapping Gaussian distributions: a left-centred distribution that constituted latent state A (μ_A_ = -6, σ = 5) and a right-centred distribution that constituted latent state B (μ_B_ = 6, σ = 5). Which state was active evolved according to a hazard rate that determined the probability of a state change after each trial. The hazard rate in a block was either 0.08 (low volatility) or 0.12 (high volatility). The low and high volatility blocks were presented in alternating order, and which condition came first was counterbalanced across participants. Participants were informed about the volatility condition of the upcoming block. Each trial consisted of an inter-stimulus-interval with a variable duration sampled from a monotonically decreasing distribution ranging from five to ten seconds (24.3 to 48.6 movable x-units), stimulus onset (“wall spawn”), and a reaction period of 1.71 seconds (8.31 movable x-units) between wall spawn and wall passage. After each trial, participants were repositioned to the central position. Task timings were set such that participants were encouraged to position themselves along the horizontal plane prior to stimulus onset to successfully pass through the gap (anticipatory movement to either the left or the right side of the central position prior to wall spawn). Auditory feedback was implemented to indicate successful passage or collision with the wall. Participants completed a three-minute practice block before starting the experiment.

### Eye data acquisition

Eye data during passive change-point oddball (Experiments 1-3) were obtained with an Eyelink 1000 device (SR Research, Osgoode, Ontario, Canada) at 1000 Hz with an average spatial resolution of 15 to 30 min arc. Eye data during active change-point navigation task in virtual reality (experiment 3), were recorded using the Pupil Labs Neon XR eye tracker mounted in the Meta Quest 3 headset at 200 Hz. Eye data during virtual reality are not presented in the current report.

### EEG data acquisition

Continuous EEG was recorded during change-point oddball in Experiment 3, from 16 scalp electrodes using a BioSemi Active Three system with a sample rate of 1024 Hz. The EEG electrodes were distributed across the scalp according to the 10–20 international system and applied using an elastic electrode cap (BioSemi). Additionally, four electro-oculogram electrodes measured eye movements and blinks: two above and below the dominant eye (vEOG) and two at the outer canthus of each eye (hEOG); each referenced to their counterpart. EEG data are not presented here but will be the scope of another report.

### Analysis of pupil responses

#### Preprocessing

Periods of blinks and saccades were detected using the manufacturer’s standard algorithms with default settings. The remaining data analyses were performed using custom-made Python software. We applied to each pupil recording (i) linear interpolation of values measured just before and after each identified blink (interpolation time window, from 200 ms before until 200 ms after blink), (ii) low-pass filtering (third-order Butterworth, cutoff, 10 Hz), (iii) removal of pupil responses to blinks and saccades by means of multiple linear regression based on double-gamma functions [125], (iv) conversion to units of modulation (percent signal change) around the mean of the pupil time series from each sequence.

#### Quantification of pupil measures

We computed the baseline pupil size for each stimulus as the mean pupil size across all values in the window -0.25 to 0 s from stimulus onset. We computed stimulus-evoked pupil responses using the following approach. First, we took the first temporal derivative by taking the discrete difference between consecutive time points and multiplying it by 100. Then we used linear projection of each single stimulus pupil time course onto the average pupil time course of that participant, to obtain a scalar measure of the overall pupil modulation amplitude (positive or negative) for each stimulus [83]:

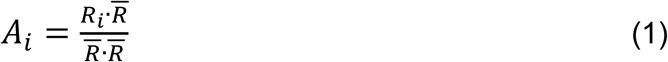

where *R_i_* is the single stimulus-evoked pupil time course, *i* indexes the stimuli, and *R* is the average evoked pupil time course of a given participant. Pupil response amplitude measures were computed for the time window 0 s to 1 s from stimulus onset, which consistently captured the full modulation participants’ pupil responses (Fig. 2B). For experiment 2, the linear projection kernel was computed as the grand average pupil time course (over all participants; due limited data available for each participant). During auditory change-point oddball, pupil response amplitudes quantified this way were not related to baseline pupil size (**Fig. S1E**,**F**), consistent with previous reports that derivative-based pupil measures show reduced baseline dependence [61,82,83].

We also considered a more conventional method of quantification, based on taking the participant-wise average of the derivative within a defined time window. This time window was chosen using a cluster-based paired permutation test of the overall mean derivative response against zero (**Fig. S2G**, **Fig. S3E**). Resulting scalar values therefore represented the mean response during a dilation or a constriction phase (**Fig. S2G-I**, **Fig. S3E-I**). Again, variations in these pupil measurements due baseline pupil size were removed via linear regression accordingly (**Fig. S1E-H**).

### Analysis approaches

#### Stimulus exclusion criteria

Change-point oddball: In all analyses, we removed the first 25 stimuli (37.5 seconds) due to heightened baseline pupil size (**Fig. S1B**, **Fig. S3A**) and for letting participants settle into the protocol. Additionally, we removed stimuli for which we measured blinks in the time window 0 s to 1 s from stimulus onset. Most stimuli remained in our analyses (**Fig. S1C**, **Fig. S3B**) and blinks were not more prevalent during state-specific oddballs vs state-specific standards (**Fig. S1D**, **Fig. S3C**).

Change-point navigation: In all task analyses, we removed the first 5 trials (33.5– 58.5 seconds) per block to let participants settle into the protocol.

#### Early and late stimuli/sequences split

Change-point oddball: When splitting data in early and late stimuli in sequences, stimulus 26 (first 25 stimuli were removed) to stimulus 300 were considered as early and 301 to 600 were considered late (**Fig. S2M**,**O**, **Fig. S4B**). When splitting data in early and late sequences, sequences 1 to 3 were considered early and sequences 4 to 6 were considered late (**Fig. S2N**,**P**, **Fig. S4C**).

#### Quantification of navigation behaviour

Change-point navigation: We computed the pre-wall position on each trial as the mean x-position in the window -0.2 to 0 s prior to wall spawn. (Fig. 5E,**F**,**G**,**H, Fig. S5A**,**C**,**D**,**E**). We interpreted this scalar as a behavioural read-out of a participant’s prior belief on each trial.

#### Stimuli after state changes

Change-point oddball: For the analysis of pupil responses after a state change (Fig. 2D, Fig. 4B, **Fig. S2F**,**K**,**O**,**P**), we considered all standard stimuli up to six stimuli after a state change, if they were not interrupted by an oddball. All standard stimuli after the sixth position were considered for the average standard response independent of oddball interruption.

Change-point navigation: For the visualisation of behaviour with respect to state changes (Fig. 5E), we considered only those state changes for which the previous state lasted at least five trials. For the analysis of pre-wall position with respect to state changes (Fig. 5F,**G**, **Fig. S5A**,**C**,**D**), we considered all trials after the sixth position for the average response.

#### Exponential fit

For all analyses with respect to state changes (Fig. 2D, Fig. 4B, Fig. 5F,**G**, **Fig. S2K**,**O**,**P**, **Fig. S5A**,**C**,**D**), we used an exponential fit to extract the time constant of decay, *τ.* The fit was implemented using the curve_fit() function from the SciPy optimize package. We considered a new steady state to be reached (in both pupil response amplitude and pre-wall position) after 5*τ* additional stimuli/trials.

#### Bayesian belief updating model

We adapted a normative model of belief updating (Fig. 4A) from previous work regarding Bayesian inference during active belief updating task or change-point detection task in humans [38,45]. In this model the perceived evidence (*LLR_n_*) at stimulus position *n*, is expressed as a log-likelihood ratio between the relative probability for stimulus 1 (*p*(*x_n_*|*S*_2_)) vs stimulus 2 (*p*(*x_n_*|*S*_2_)). The evidence is added to a preexisting prior belief (**Ψ***_n_*) upon stimulus encounter to form the posterior belief (*L_n_*) for this stimulus. Thus, the evidence and the posterior is defined as:

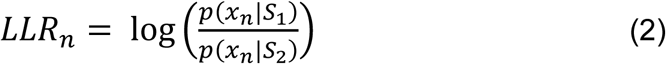

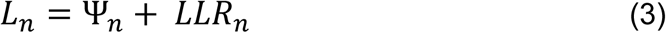

This posterior is transformed in a non-linear manner to act as the new prior for the next stimulus encounter. This non-linear transformation is guided by the “hazard rate” (*H*), which can be seen as the model’s estimation of the latent state volatility. A low hazard rate assumes a stable world with no state changes, while a higher hazard rate discounts the influence of the posterior belief on the prior of the next stimulus encounter, making the system more sensitive to detect change-points. The hazard rate-based transformation of posterior beliefs to the next prior is expressed by:

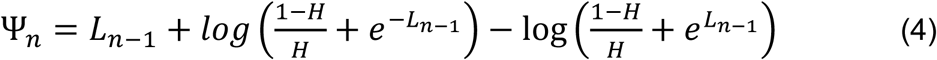

Stimulus-wise change-point probability (*CPP_n_*) can be derived as follows [45,89]:

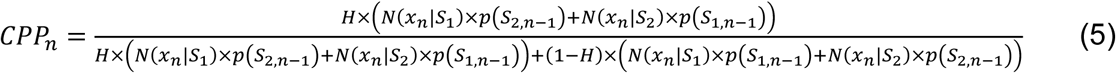

Here, *N*(*x_n_*|*S*_1/2_) denotes the probability that sample *x_n_* is stimulus 1 (*S*_1_) or stimulus 2 (*S*_2_) given a normal distribution. *p*(*S*_1/2,*n*−1_) is the probability that the previous sample was stimulus 1 (*S*_1_) or stimulus 2 (*S*_2_). Of note, the presentation of either one of two discrete tones as input restricts the log-likelihood ratio values of evidence to be either a positive or negative expression of the same value.

Stimulus-by-stimulus uncertainty (*U_n_*) was defined as follows:

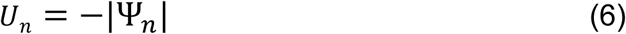

To compute stimulus-by-stimulus change-point probability and uncertainty in change-point oddball, we used the objective hazard rate *H*, except for one control analysis where we varied *H* (**Fig. S4D**). In the change-point navigation task, we used the average of the two hazard rate conditions (*H* = 0.1). Final results (Fig. 5K) are robust to using *H* = 0.08 or *H* = 0.12, corresponding to the objective hazard rate used in the low or high hazard rate condition, respectively (**Fig. S5O**,**P)**.

#### Time-wise regression analysis pupil responses

To analyse the strength of change-point probability encoding in the pupil per subject, we implemented a time-wise linear regression model as used before [45,89] on the pupil response time course (Fig. 4C, **5J,K**):

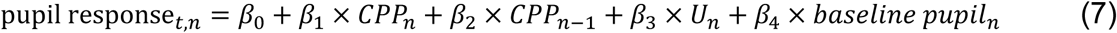

Where *t* indicates time point, *n* indicates stimulus position in the sequence, *U_n_* is the uncertainty (inverse of absolute prior belief), other variables are defined as before, and *β*_0_, *β*_1_, *β*_2_, *β*_3_, and *β*_4_ are the parameters of the fit. Note that we did not include |*LLR_n_*| (as in [45,89]), since discrete stimulus identities in change-point oddball render this variable to fixed values. Predictors were z-scored prior to fitting.

#### Simulations virtual players change-point navigation task

We simulated the pre-wall positions of four synthetic agents (“players”) on the change-point navigation task. The *optimal player* updates its behaviour according to the Bayesian belief updating model as described above. We used a hazard rate of 0.1 (H = 0.1) to account for the collapsing across hazard rate conditions of 0.08 and 0.12. The *perfect accumulator* updates its behaviour based on full trial history. We simulated this by setting H = 0.0001, which essentially linearizes the non-linear transform function and gives it a slope of 1, meaning no loss of information from previous posterior to next prior. Both the optimal player and perfect accumulator predictions were scaled to [μ_A_, μ_B_] such that their pre-wall positions mapped onto task-relevant spatial dimensions. The *last sample player* does not accumulate evidence but bases its current belief solely on the last trial. We simulated this by setting its behavioural prior to μ_A_ = -6 or μ_B_ = 6 if the wall opening on the previous trial was on the left or right, respectively. Lastly, we simulated a *middle dweller* that did not implement pre-sorting but always remained at the central position (x = 0) prior to wall spawn.

#### Strategy regression pre-wall position behaviour

To evaluate the usage of different strategies in the change-point navigation task, we calculated pre-wall positions of the different synthetic agents and used those to predict each participant’s pre-wall position (Fig. 5I):

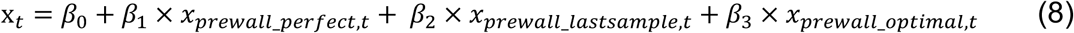

Here, *t* indicates trial number, x*_t_* is pre-wall position of a subject on trial *t*, anx*_prewall_perfect,t,_* x*_prewall_lastsample,t_* and x*_prewall_optimal,t_* are the pre-wall positions of the following synthetic agents: the perfect accumulator, last sample player, and optimal Bayesian player, respectively. *β*_0_, *β*_1_, *β*_2_ and *β*_3_ are the parameters of the fit. The intercept accounts for any consistent bias, such as “middle dweller” behaviour. Predictors were z-scored prior to fitting.

#### Regression behavioural belief update

To characterize the influence of prediction errors, change-point probability and uncertainty on each participant’s belief updates, we used a linear regression model as implemented previously in [50,90] (**Fig. S5F**):

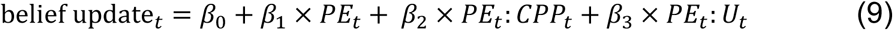

Here, belief update at trial *t* is defined as the difference between pre-wall position on trial *t* + 1 and trial *t*, *PE* (prediction error) on trial *t* is defined as the difference between prewall position on trial *t* and gap position on trial *t*, other variables are defined as before, and *β*_0_, *β*_1_, *β*_2_ and *β*_3_ are the parameters of the fit. Predictors were z-scored prior to fitting.

#### Regression analysis pupil response scalars

To summarize the strength of change-point probability encoding in the pupil per subject, we implemented subject-wise linear regression models on the pupil response scalar:

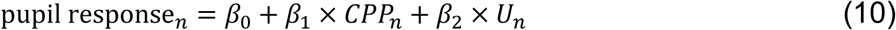

Where *n* indicates stimulus position in the sequence, and *β*_0_, *β*_1_ and *β*_2_ are the parameters of the fit. *β*_1_ was used as the dependent variable in the below regression model (Eq. 11) linking the two tasks (*βCPP*). Predictors were z-scored prior to fitting.

#### Linking pupil responses in change-point oddball and behaviour in the change-point navigation task

To test if the magnitude of the influence of *PE*, *CPP* and *U* on belief updates in the change-point navigation task predicts change-point probability encoding in the pupil in change-point oddball, we extracted the regression coefficients of the subject-wise regression above (Eq. 10) (*βPE*, *βPE*: *CPP*, *βPE*: *U*) and used these to predict the change-point probability regression coefficients for each participant from Eq. 10 (*βCPP*) (Fig. 5K, **Fig. S5M-R**).

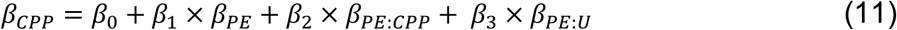

Predictors were z-scored prior to fitting.

### Statistical comparisons

We used non-parametric paired permutation tests to assess statistical significance. All statistical comparisons were performed across participants, using the mean per participant as observation. For each comparison, we randomly permuted the labels of the observations within a participant and recalculated the difference between the two group means (10K permutations). The p-value associated with the original difference between the means was given by the fraction of shuffles in which the original difference was exceeded by the difference between the means obtained for the shuffled data.

For the non-parametric paired permutation test of the interaction between state and stimulus identity (Fig. 2C, **Fig. S2H**-**J**, Fig. 3B, **Fig. S3F**,**G**), we generated a difference of differences vector according to:

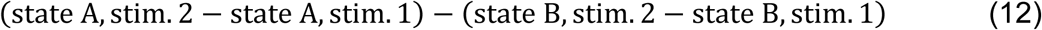

The resulting vector was then compared with a zero-vector using a non-parametric paired permutation test (**Fig. S2C**). To assess the strength of this interaction (F-Value, p-value) we performed a repeated measure ANOVA with the pupil scalar as dependent variable and state and stimulus as within subject factors. The data were aggregated using the mean (**Fig. S2F**).

To test the correlation values between pupil scalar and change-point probability, we calculated the correlation (Pearson’s correlation) between the binned pupil scalar and change-point probability per participant and compared the resulting r-values with a zero-vector using a non-parametric paired permutation test (Fig. 4D, **Fig. S4B**,**C**). In the case of early-late comparisons, r-value vectors were compared with each other using the same test (**Fig. S4B**,**C)**.

To test significance of the regression coefficients from the time-wise regression analysis of pupil responses (Fig. 4C, Fig. 5J,**K**), we compared the participant-wise beta-vectors at each time point with a zero-vector using a non-parametric one-sample cluster-based permutation test. This test identifies clusters of adjacent time points showing consistent non-zero regression coefficients across participants and evaluates their significance against a permutation-derived null distribution. Clusters-level p-values were used to determine significance.

For the non-parametric paired permutation tests between regression coefficients of different synthetic players in the change-point navigation task (Fig. 5I), we compared the beta-vectors of the different players with one another, and each player with a zero-vector.

#### Bootstrapping

For every bootstrapping approach the bootstrap was performed 10K times with replacement (Fig. 2D, **Fig. S2K**,**O**,**P**, Fig. 5F, **Fig. S5A**,**B**,**G**,**H**,**K**,**L**, Fig. 5K, **Fig. S5M-R**). For exponential fit bootstraps (Fig. 2D, **Fig. S2K**,**O**,**P**, Fig. 5F, **Fig. S5A**), data was bootstrapped grouped by participant and stimulus position after state change. The resulting boots were used to create an exponential fit distribution, which was displayed as a 95% confidence interval in the exponential decay figures (Fig. 2D, **Fig. S2K**,**O**,**P**, Fig. 5F, **Fig. S5A**).

The 95% bootstrap confidence intervals of tau estimates from the exponential fits were displayed as error bars around individual participant’s tau estimates (**Fig. S5B**). Similarly, the 95% confidence intervals resulting from bootstrapping of regression analyses were displayed as error bars around individual participant’s regression coefficients in the figures (Fig. 5K, **Fig. S5M-R**).

## Data, Materials and Software availability

Data and analysis scripts will be made publicly available upon publication.

## Acknowledgements

We are grateful to all participants. We thank Yihsin Chuang and Noell Vermeer for their help with data collection in experiment 2.

## Author contributions

Conceptualization, LK, JWdG; Investigation, LK, JMH, EvG, JVT, MG, MS; Formal analysis, LK, JMH, EvG, JWdG; Resources, SM, PM, SvG; Writing— original draft, LK, JMH, JWdG; Writing—review and editing, LK, JMH, PM, SvG, MS, JWdG; Supervision, JWdG; Funding acquisition, MS, JWdG.

## Supplementary figures

**Suppl. Fig. 1.**
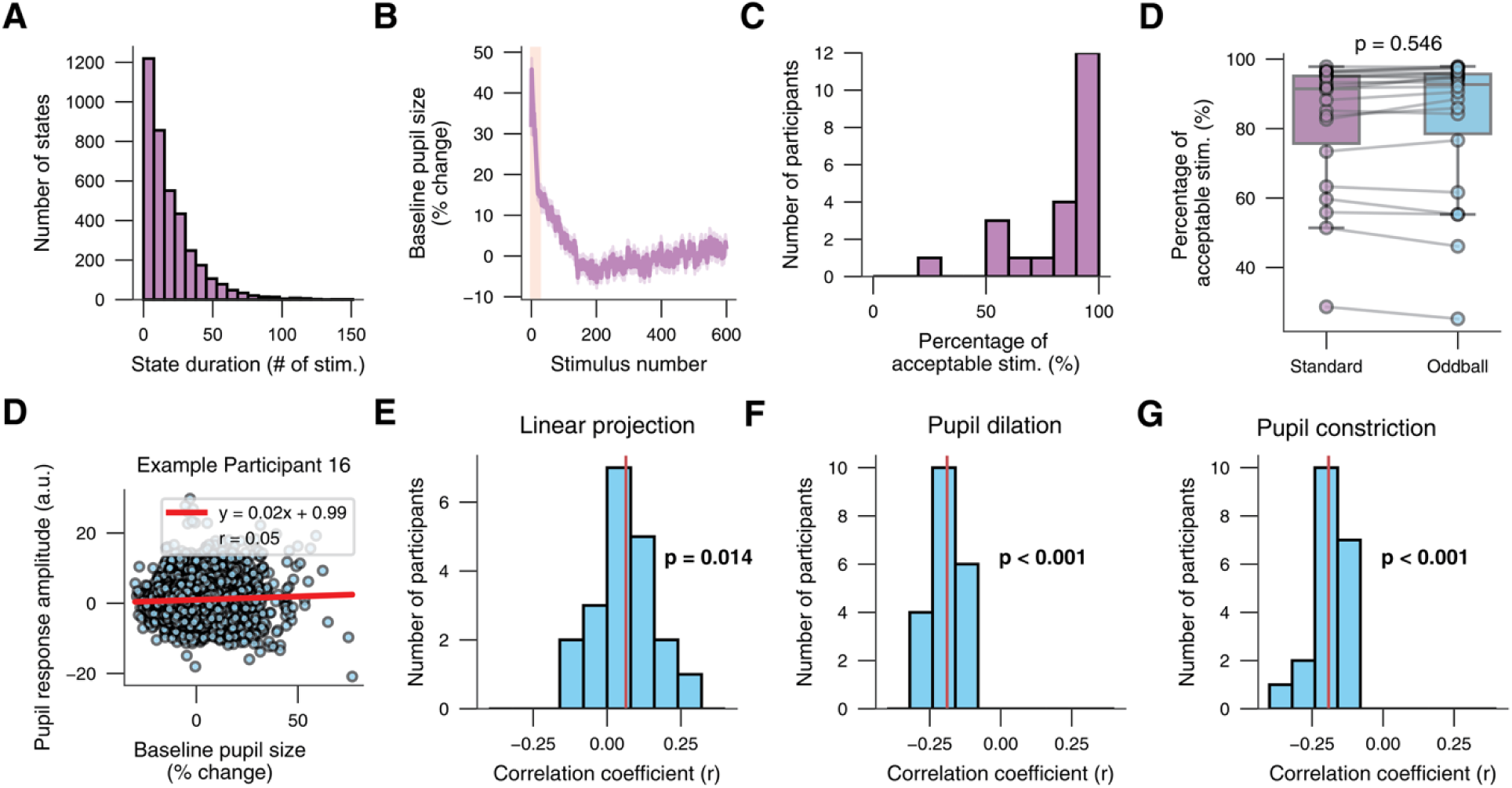
**(A)** Histogram of state durations (i.e. number of stimuli between two consecutive state changes) in the change-point oddball paradigm with a hazard rate of 0.05. **(B)** Baseline pupil size against stimulus numbers since recording start, red bar marks first 25 stimuli, which were excluded. **(C)** Distribution of participant-wise percentage of acceptable stimuli (% stimulus periods without recorded blink). **(D)** Comparison of percentage of acceptable standard and oddball stimuli, dots indicate participants. Stats, nonparametric paired permutation test. **(E)** Pupil response amplitude against baseline pupil size of all stimuli for example participant 16. Linear regression equation (red) and Pearson correlation coefficient (r) in legend. **(F)** Distribution of participant-wise Pearson correlation coefficients of linear projection values, mean correlation coefficient (r) in red. Stats, non-parametric paired permutation test of r-values against zero vector. **(G)** Same as F for pupil dilation values. **(H)** Same as G and F for pupil constriction values. Panel A: shading SEM across participants (N = 20, experiment 1).

**Suppl. Fig. 2.**
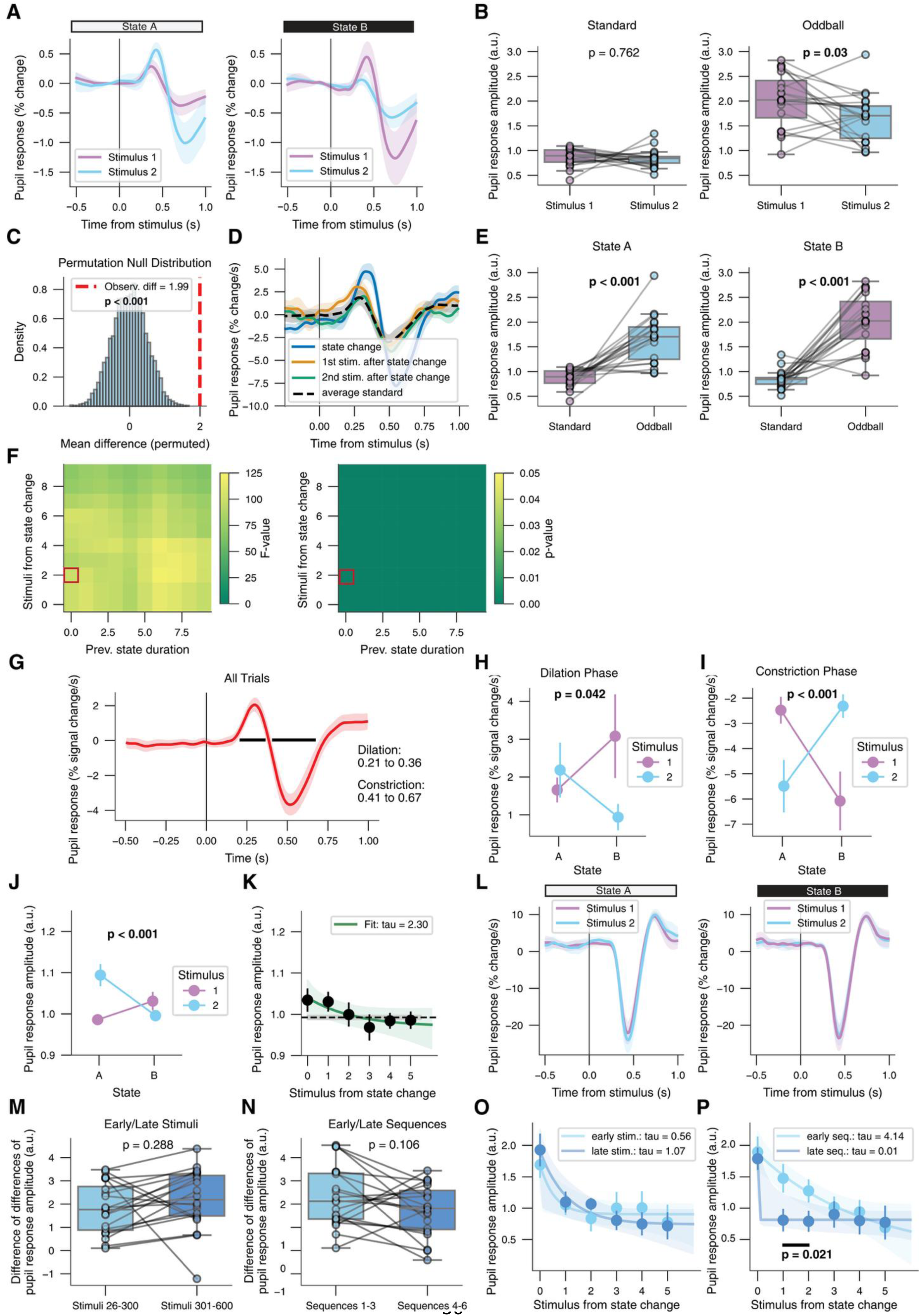
**(A)** Pupil response (non-derivative) locked to each stimulus, separately for states and stimuli. Black line, stimulus onset. **(B)** Comparison between pupil response amplitudes towards stimulus 1 and stimulus 2, separated by standards or oddballs to show stimulus specificity. Dots indicate participants. Stats, non-parametric paired permutation test. **(C)** Permutated null distribution of difference of difference ((state A, stim. 2 state A, stim. 1) (state B, stim. 2 state B, stim. 1)) vs zero vector. Red dotted line indicates observed mean difference. Stats, non-parametric paired permutation test of difference of differences against zero vector. **(D)** Pupil response (first derivative) time-locked to state-specific standard stimulus onset, separately for standard stimulus position after state change and average standard stimulus response (>6 stimulus after state change). **(E)** Comparison between pupil response amplitudes towards standards and oddballs, separated by state, dots indicate participants. Stats, non-parametric paired permutation test. **(F)** Heatmap of repeated measure ANOVA F-values (upper) and corresponding p-values (lower) exploring the effect of stimulus and state on pupil response amplitude for different selection criteria (stimulus from state change and previous state duration), F-values and p-values indicate a significant interaction at all configurations, red square indicates the criteria we used in our analysis. Stats, for stimuli after state change >= 2 and prev. state duration >= 0, RM ANOVA: main effect stimulus: F(1,19) = 3.49, p = 0.077; main effect state: F(1,19) = 6.46, p = 0.02; stimulus x state interaction: F(1,19) = 97.87, p < 0.001. **(G)** Average pupil response (first derivative) of all considered stimuli. Black bars, windows of significant difference from zero. Stats, cluster-based paired permutation test against zero vector. **(H)** Average pupil response (% signal change) interaction between stimulus and state. Dilation phase for averaging based on timeframes defined by G. Stats, non-parametric paired permutation test of difference of differences against zero vector. **(I)** Same as H but for constriction phase as defined by G. **(J-L)** Pupil responses in visual change-point oddball (Experiment 1). **(J)** Pupil response amplitude (Methods) interaction between stimulus and state. Note that the dynamic range of the linearly projected pupil response amplitudes is more limited during the visual sequences than during the auditory sequences (**Fig. 2C,D**). This is because the kernel used for linear projection combines a contrast-related constriction component with a putative arousal-related dilation component, whereas deviance detection would be expected to influence only the latter. Stats, non-parametric paired permutation test of difference of differences against zero vector. **(K)** Pupil response amplitude for state-specific standard stimuli locked to state changes (uninterrupted by oddballs). Green line, exponential decay fit; green shading, 95% bootstrapped confidence interval of that fit; dashed black line, pupil response amplitude to state-specific standards across all stimuli >6 from state change. Stats, cluster-based paired permutation test between data points and dashed line. **(L)** Pupil response (first derivative) locked to each stimulus, separately for states and stimuli. Black line, stimulus onset. **(M)** Comparison between difference of differences (state-stimulus interaction) of early and late stimuli in a sequence (Methods), dots represent participants. Stats, non-parametric paired permutation test. **(N)** Same as M for comparison of early and late sequences (Methods). **(O)** Pupil response amplitude for state-specific standard stimuli locked to state changes (uninterrupted by oddballs), split by the early and late stimuli in sequences, light blue (early stimuli) or dark blue (late stimuli) line, exponential decay fit (including 95% bootstrapped confidence interval shown as shading). Stats, cluster-based paired permutation between early and late. **(O)** Same as N for comparison of early (light blue) and late (dark blue) sequences. Panel, A, B, E, H, I, J, L, M, N: as in Fig. 2B,C, all but first two stimuli in each state. Panel F, K, O, P: as in Fig. 2D, first six uninterrupted standard stimuli of each state (D only shows first three). Panel A, D, G, H, I, J, L: shading and error bars, SEM across participants (N=20, experiment 1).

**Suppl. Fig. 3.**
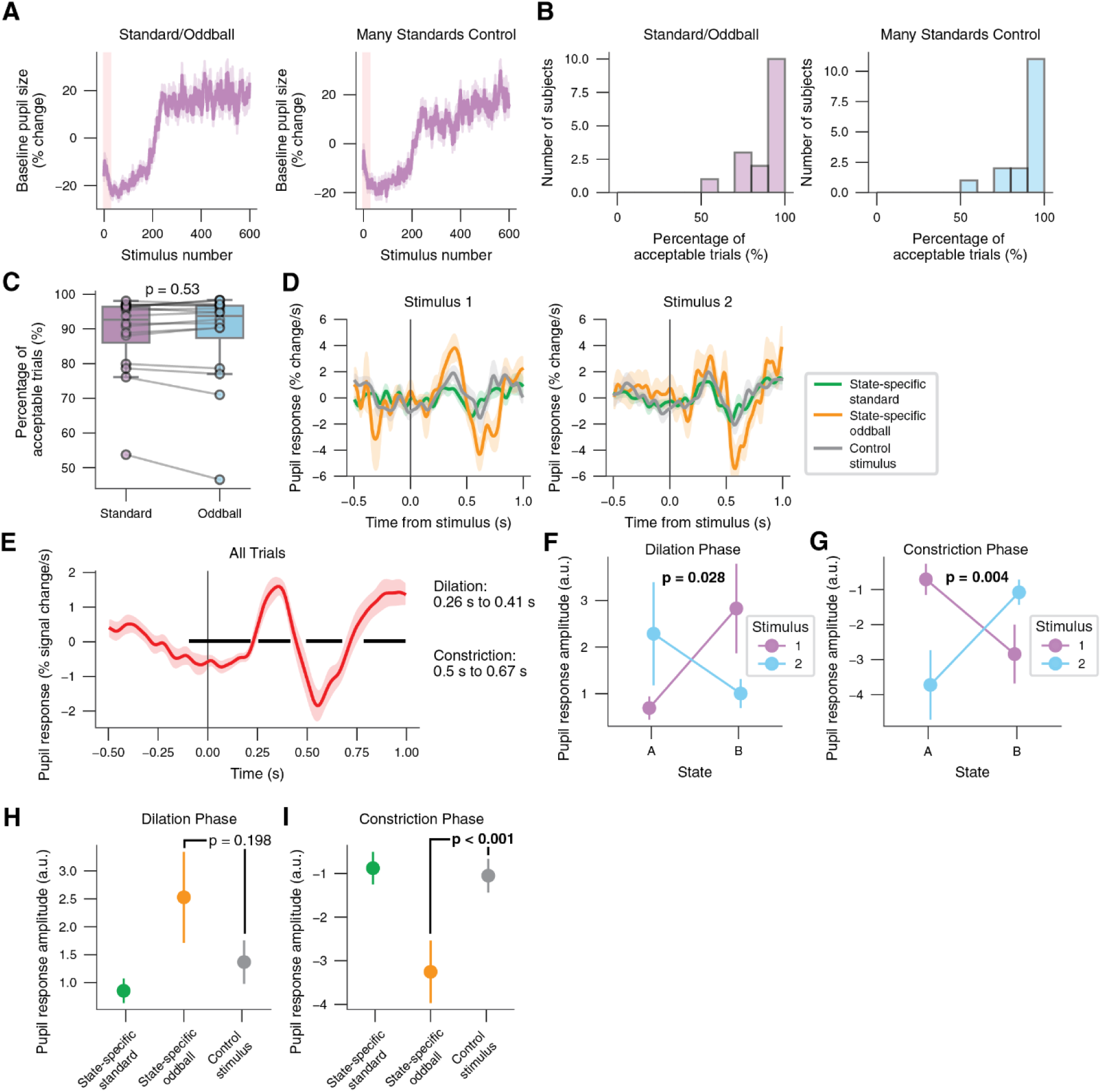
**(A)** Baseline pupil size against stimulus number since recording start for change-point oddball and many standards control protocol, note the elevated pupil response toward the change in screen brightness around mid-sequence. (**B)** Distribution of participant-wise percentage of acceptable stimuli (% stimulus periods without recorded blink) for change-point oddball and many standards control protocol. **(C)** Comparison of percentage of acceptable standard and oddball stimuli (% stimuli periods without recorded blink) during change-point oddball. Dots indicate participants. Stats, non-parametric paired permutation test. **(D)** Pupil response (first derivative) time-locked to each stimulus, separately for stimulus and context, black line indicates sound onset. **(E)** Average pupil response (first derivative) of all considered stimuli (many standard control and change-point oddball) time-locked to stimulus onset. Black bars, windows of significant difference from zero. Stats, cluster-based one-sample permutation test against zero, here only dilation and constriction phases are considered. **(F)** Average pupil response (% signal change) interaction between stimulus and state identity. Dilation phase for averaging based on timeframes defined by E. Stats, nonparametric paired permutation test of difference of differences against zero vector. **(G)** Same as F but for constriction phase as defined by E. **(H)** Comparison of stimulus evoked pupil responses in standard (green), oddball (orange) and control (dark grey) context, for dilation phase defined by E. Stats, non-parametric paired permutation test. **(I)** Same as H but for constriction phase as defined by E. Panel D, F, G. H, I: as in Fig. 2 B,C, all but first two stimuli in each state. Panel A, D, E, F, G, H, I: shading and error bars, SEM across participants (N = 15, experiment 2).

**Suppl. Fig. 4.**
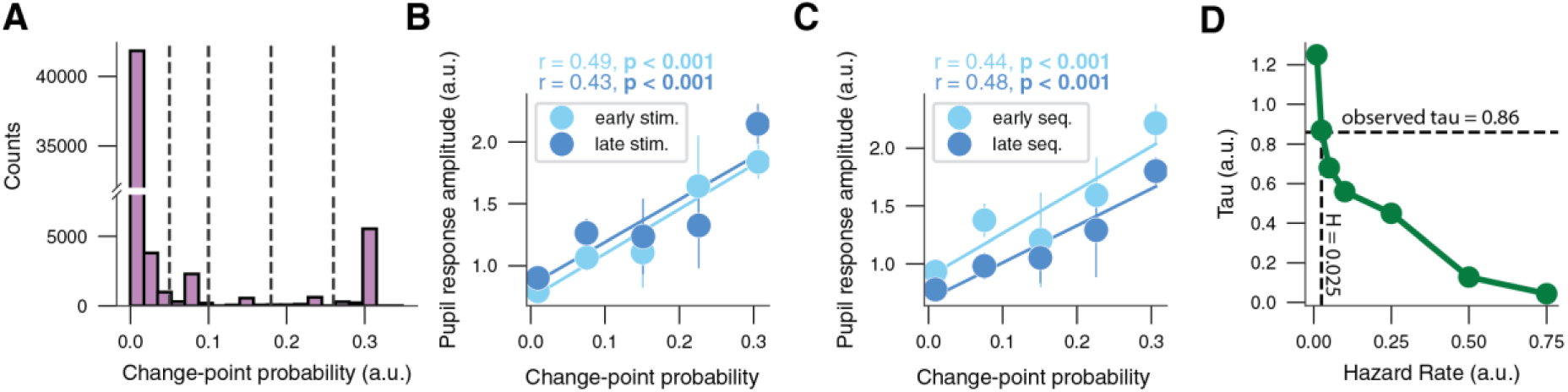
**(A)** Calculated change-point probability estimates. Dashed line, cutoffs for bins at 0, 0.05, 0.1, 0.18, 0.26, 1. **(B)** Correlation between pupil response amplitude and change-point probability split by early (light blue) and late (dark blue) stimuli in sequences (Methods). Change-point probability values binned according to A. Stats, Pearson’s correlation (r) per participant and non-parametric paired permutation test of participant-wise r-values against zero vector. No significant difference between the correlation for early vs, late stimuli (non-parametric paired permutation test, mean difference = -0.05, p = 0.59). **(C)** Same as B but split for early (light blue) and late (dark blue) sequences (Methods). No significant difference between the correlation for early vs. late sequences (non-parametric paired permutation test, mean difference = -0.04, p = 0.701). **(D)** Tau values of modelled change-point probability after state change responses (as in Fig. 4B for different subjective hazard rates). Panels B,C: error bars, SEM across participants (N = 20; experiment 1).

**Suppl. Fig. 5.**
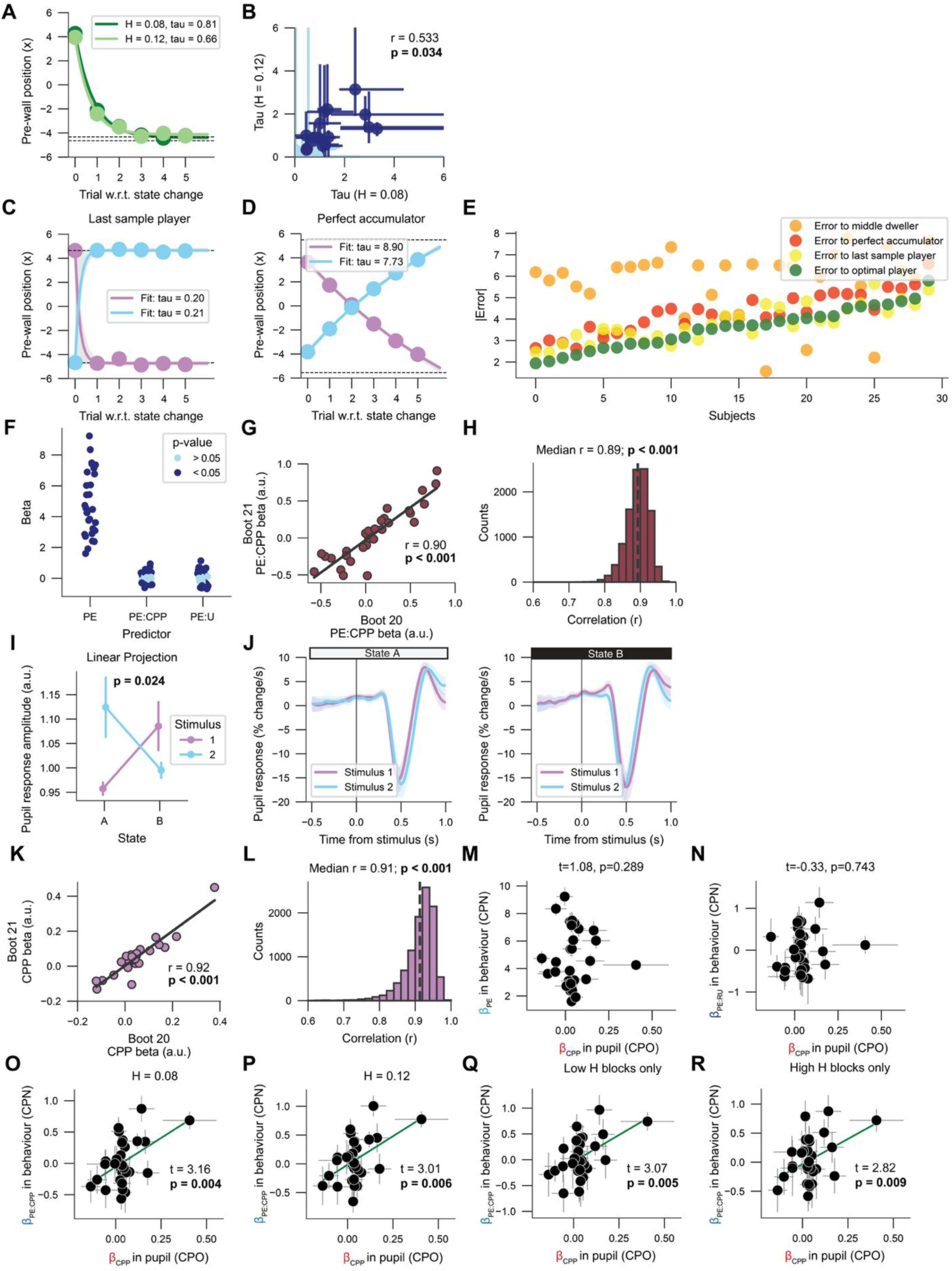
**(A)** Pre-wall position locked to state changes, separately for the two hazard rates. Green and light green lines, exponential decay fit of H = 0.08 and H = 0.12, respectively; light grey shading, 95% bootstrapped confidence interval of fits; dashed black lines, pre-wall position across all stimuli >5 from state change. No significant difference between the two hazard rate conditions (non-parametric paired permutation test, p = 0.47). **(B)** Correlation of individual participants’ fitted tau estimates for the two hazard rate conditions; horizontal and vertical error bars, 95% bootstrapped confidence interval of fits of low and high hazard rate conditions, respectively. Light blue, participants for whom at least 5% of bootstrapped tau fits reached lower or upper boundary in either condition, i.e. poorly identified tau fits; blue, participants with well-identified tau fits. **(C)** Last sample player pre-wall position locked to state changes (H = 0.1 used to simulate data) (Methods). Pink and blue lines, exponential decay fit of state A and B, respectively; dashed black lines, pre-wall position across all stimuli >5 from state change. **(D)** Perfect accumulator pre-wall position locked to state changes (H = 0.1 used to simulate data) (Methods). Pink and blue lines, exponential decay fit of state A and B, respectively; dashed black lines, pre-wall position across all stimuli >5 from state change. **(E)** Absolute error between participants’ pre-wall position and simulated pre-wall positions of synthetic agents. Subjects sorted by absolute error relative to the optimal player. Majority of subjects exhibit behaviour that best reflects the optimal strategy. **(F)** Subject-wise regression coefficients of the regression model belief update ∼ PE + PE:CPP + PE:U (Eq. 9). Dots indicate participants. **(G**,**H)** Bootstrapping reliability of individual differences in PE:CPP betas in the change-point navigation task. **(G)** PE:CPP regression coefficients for neighbouring example boots plotted against one another. Every data point is a participant. Stats, Pearsons’s correlation. **(H)** Distribution of correlation coefficients of PE:CPP betas (as in panel G) from all neighbouring boots. Dotted black line, r = 0.5 mark, a commonly used threshold for acceptable reliability. P-value reflects the fraction of correlation coefficients being smaller than 0.5. Repeating this analysis with Spearman rank-based correlation yielded a median correlation coefficient of r = 0.89, p < 0.001. **(I)** Pupil response (first derivative) locked to each stimulus in audio-visual change-point oddball (Experiment 3), separately for states and stimuli. Black line, stimulus onset. **(J)** Pupil response amplitude interaction between stimulus and state in audio-visual change-point oddball (Experiment 3). Stats, non-parametric paired permutation test of difference of differences against zero vector. **(K**,**L)** Bootstrapping reliability of individual differences in CPP betas in change-point oddball. **(K)** CPP regression coefficients for neighbouring example boots plotted against one another. Every data point is a participant. Stats, Pearson’s correlation. **(L)** Distribution of correlation coefficients of CPP betas (as in panel I) from all neighbouring boots. Repeating this analysis with Spearman rank-based correlation yielded a median correlation coefficient of r = 0.79, p < 0.001. **(M**,**N)** Linear regression model fit for the two other predictors in the regression model βCPP ∼ βPE + βPE*CPP + βPE*U (Eq. 11). **(M)** Regression coefficients of PE in behaviour (βPE) vs. regression coefficients of CPP in pupil (βCPP). **(N)** Regression coefficients of PE:U in behaviour (βPE) vs. regression coefficients of CPP in pupil (βCPP). **(O**, **P)** Regression coefficients from the same regression model but with model variables (CPP, U) computed using H = 0.08 and H = 0.12, corresponding to the objective hazard rates in the low and high hazard rate conditions, respectively. **(O)** Regression coefficients of PE:CPP in behaviour (βPE:CPP) significantly predict regression coefficients of CPP in pupil (βCPP) using the low hazard rate-derived computational variables. **(P)** Same as O when using the high hazard rate-derived computational variables. **(Q**, **R)** Regression coefficients from the same regression model but with model variables (CPP, U) computed using either only low or high hazard rate condition trials **(Q)** Regression coefficients of PE:CPP in behaviour (βPE:CPP) significantly predict regression coefficients of CPP in pupil (βCPP) using only trials in the low hazard rate blocks. **(R)** Same as Q when using only trials in the low hazard rate blocks. Panel B,M,N,O,P,Q,R: Error bars bootstrapped 95% confidence interval (10K bootstraps; Methods). Panel I,J: shading and error bars, SEM across participants. (N = 30; experiment 3).

